# KIT ligand protects against both light-induced and genetic photoreceptor degeneration

**DOI:** 10.1101/752030

**Authors:** Huirong Li, Lili Lian, Bo Liu, Yu Chen, Jinglei Yang, Shuhui Jian, Jiajia Zhou, Ying Xu, Xiaoyin Ma, Jia Qu, Ling Hou

## Abstract

Photoreceptor cell degeneration is a major cause of blindness and a considerable health burden during aging but effective therapeutic or preventive strategies have not so far become commercially available. Here we show in mouse models that signaling through the tyrosine kinase receptor KIT protects photoreceptor cells against both light-induced and inherited retinal degeneration. Upon light damage, photoreceptor cells upregulate Kit ligand (KITL) and activate KIT signaling, which in turn induces nuclear accumulation of the transcription factor NRF2 and stimulates the expression of the antioxidant gene *Hmox1*. Conversely, a viable *Kit* mutation promotes light-induced photoreceptor damage, which is reversed by experimental expression of *Hmox1*. Furthermore, overexpression of KITL from a viral AAV8 vector prevents photoreceptor cell death and partially restores retinal function after light damage or in genetic models of human retinitis pigmentosa. Hence, application of KITL may provide a novel therapeutic avenue for prevention or treatment of retinal degenerative diseases.

## INTRODUCTION

Retinal degeneration due to photoreceptor loss is a major threat to human health as progressive vision loss severely interferes with a person’s daily activities. Such photoreceptor cell loss is common to a number of degenerative eye disorders including cone dystrophy, retinitis pigmentosa (RP) and the highly prevalent age-related macular degeneration (AMD) (*1–3*). Each of these disorders is influenced, to various degrees, by mutations in a number of distinct genes. RP, for instance, has been associated with mutations in~80 genes and can be inherited as an autosomal-dominant, autosomal-recessive, or X-linked trait (*4*). It is characterized initially by degeneration of rod photoreceptors, followed by a loss of cone photoreceptors and of the photoreceptor-trophic retinal pigment epithelium (RPE). Consequently, patients suffer from progressive night blindness, tunnel vision and rarely may become totally blind (*4, 5*).

Currently, one of the major challenges for the field of retinal degenerative diseases is to provide strategies to prevent or delay the onset of photoreceptor cell loss. Several approaches are currently under investigation to treat vision loss in patients suffering from retinal degenerations, including retinal prostheses implants, stem cell transplants, and gene therapy (*6, 7*). Although gene therapy has recently been approved for a specific causative mutation (RPE65), there are still no broadly applicable and efficacious treatments available to prevent or cure vision loss caused by photoreceptor cell degeneration. Alternatively, photoreceptor cell death could be prevented by strengthening endogenous prosurvival mechanisms or by directly blocking cell death (*9, 10*). In fact, several neuroprotective factors have been identified based on results obtained with transgenic and conditional knockout animals and pharmacological treatments (*11, 12*). They include ciliary neurotrophic factor (CNTF), glial cell–derived neurotrophic factor (GDNF), pigment epithelium-derived factor (PEDF) and rod-derived cone viability factor (RdCVF), all of which might potentially be used to treat photoreceptor degeneration in humans (*13–17*). Their clinical application, however, is hampered by relatively short half-lives, systemic side effects and reduced efficacy, and none are commercially available for clinical use (*7, 10*). Therefore, there is a need to identify additional factors that are capable of improving photoreceptor survival and that may have pharmacokinetic properties that differ from those already studied.

Retinal degeneration, regardless of whether it is due to intrinsic genetic or extrinsic environmental conditions, is normally associated with an induction of damaging effector molecules. As a corollary of damage, the eye may respond with the induction of damage-limiting, endogenous neuroprotective factors such as the ones mentioned above (*18–20*). This fact prompted us to search for additional neuroprotective factors after experimental light damage (LD). It is known that limited light stress activates the FGF2/gp130 signaling pathway (*11, 21*) and that prolonged exposure to light up-regulates CNTF and FGF2, although usually at levels too low to protect against retinal degeneration (*11, 22, 23*). That LD is a valid strategy to identify novel retinoprotective factors is underscored by the role excessive light may play in worsening genetically influenced retinal degenerations such as AMD (*24–26*).

To discover novel retinoprotective factors, we applied an RNA-seq analysis to mouse retinas after prolonged LD. This approach confirmed the induction of a number of genes previously found to be altered during LD (*27–29*) and identified a number of novel genes not previously recognized in this context. One of them is *Kit ligand* (*Kitl*) whose protein product, KITL (also known as stem cell factor, SCF, or mast cell growth factor, MGF), stimulates an important signaling pathway by interacting with its receptor, KIT (also known as the proto-oncogene c-KIT) (*30–32*). KIT is a multi-domain transmembrane tyrosine kinase (*33*) expressed in a variety of cells including melanocytes, germ cells, cells of the hematopoietic lineage, and interstitial cells of Cajal in the gastrointestinal tract and is required for their normal development and function (*34–37*). Interaction of KIT with its ligand leads to receptor dimerization and autophosphorylation which in turn activates the MAPK pathway, phosphatidylinositol 3’-kinase (PI3K), JAK/STAT and SRC family members (*30, 31*). Gain of function mutations can lead to aggressive mastocytosis, gastrointestinal stromal tumors, melanoma and urticaria pigmentosa, and loss of function mutations can lead to piebaldism and hearing loss (*38–41*). In mice, loss-of-function mutations in *Kit* cause severe anemia, pigmentation abnormalities, sterility, mast cell deficits, lack of the interstitial cells of Cajal in the gut, spatial learning memory deficits and defects in peripheral nerveregeneration (*30, 31, 42, 43*). Nevertheless, although KIT has been found to be expressed in retinal progenitor cells (*44, 45*), its role in the adult retina is still unknown.

Here we find that mice homozygous for the *Kit*^*Wps*^ mutation (*46*) show an exacerbated photoreceptor degeneration and that overexpression of KITL can prevent light-induced photoreceptor cell death in *Kit+/+* mice. Furthermore, we show that KIT signaling stimulates the expression of *Hmox1* in an NRF2-dependent manner and that experimental expression of *Hmox1* in *Kit*^*Wps*^ homozygotes prevents light-induced photoreceptor cell degeneration. Furthermore, we show that overexpression of KITL prevents photoreceptor cell death and partially rescues the retinal dysfunction in mouse genetic models of retinitis pigmentosa. Hence, our findings suggest a mechanism by which KITL/KIT signaling contributes to protection of photoreceptor cells from degeneration and which may potentially lead to novel therapeutic strategies in retinal degenerative disorders.

## RESULTS

### Light damage upregulates endogenous KITL in the retina

Previous results have shown that light stress induces endogenous factors capable of protecting photoreceptor cells (*11, 21*). Hence, we used LD in light sensitive BALB/c albino mice to search for such inducible factors. Retinal damage is usually accompanied by reactive gliosis, characterized by the accumulation of filamentous proteins such as glial fibrillary acidic protein (GFAP) and formation of a glial scar (*47*). As shown in Supplemental Figure 1, after 1 day of continuous exposure (15,000 Lux), the retinas showed no obvious signs of retinal degeneration, but increased expression of GFAP. After three days of continuous exposure, however, there was significant retinal degeneration (Supplemental Figure 1A to C). Therefore, we used the 1-day exposure to identify novel protective genes (Fig. 1A) and longer exposures to test for effects on late-stage damage. RNA-seq data showed a large number of genes whose expression was changed in the LD-treated retinas compared with the non-treated ones (Fig. 1A), including known light-inducible genes such as *Gfap, Edn2, Fgf2, Ccl3* and *Ccl12* (Fig. 1B) and hence supporting the validity of this approach (*27–29*). A KEGG (Kyoto Encyclopedia of Genes and Genomes) analysis identified the MAPK pathway among the top 20 altered signaling pathways (Fig. 1C), with increased expression, for instance, of the genes encoding LIF, KITL and NGF (Fig. 1D). RT-PCR confirmed their differential expression identified by the RNA-seq analysis. In particular, *Kitl* expression was significantly increased after LD (Supplemental Figure 1D). Because KIT signaling plays vital roles in a variety of cell types, we focused on this pathway in the eye. Quantitation of western blot bands showed that both mature transmembrane and cleaved soluble forms of KITL (45 and 19 kDa respectively) were increased (m-KITL, 2.4 ± 0.9 fold; s-KITL, 4.5 ± 1.6 fold; n=5) (Fig. 1 E and F). Phosphorylated KIT (indicative of activate KIT), though not total KIT, was also increased (3.14 ± 0.94, n=5) (Fig. 1 G and H). Immunostaining showed that upregulation of KITL was restricted to photoreceptor inner segments (Fig. 1I). These results clearly show that following prolonged light exposure, upregulation of KITL leads to activation of KIT signaling.

**Figure 1.**
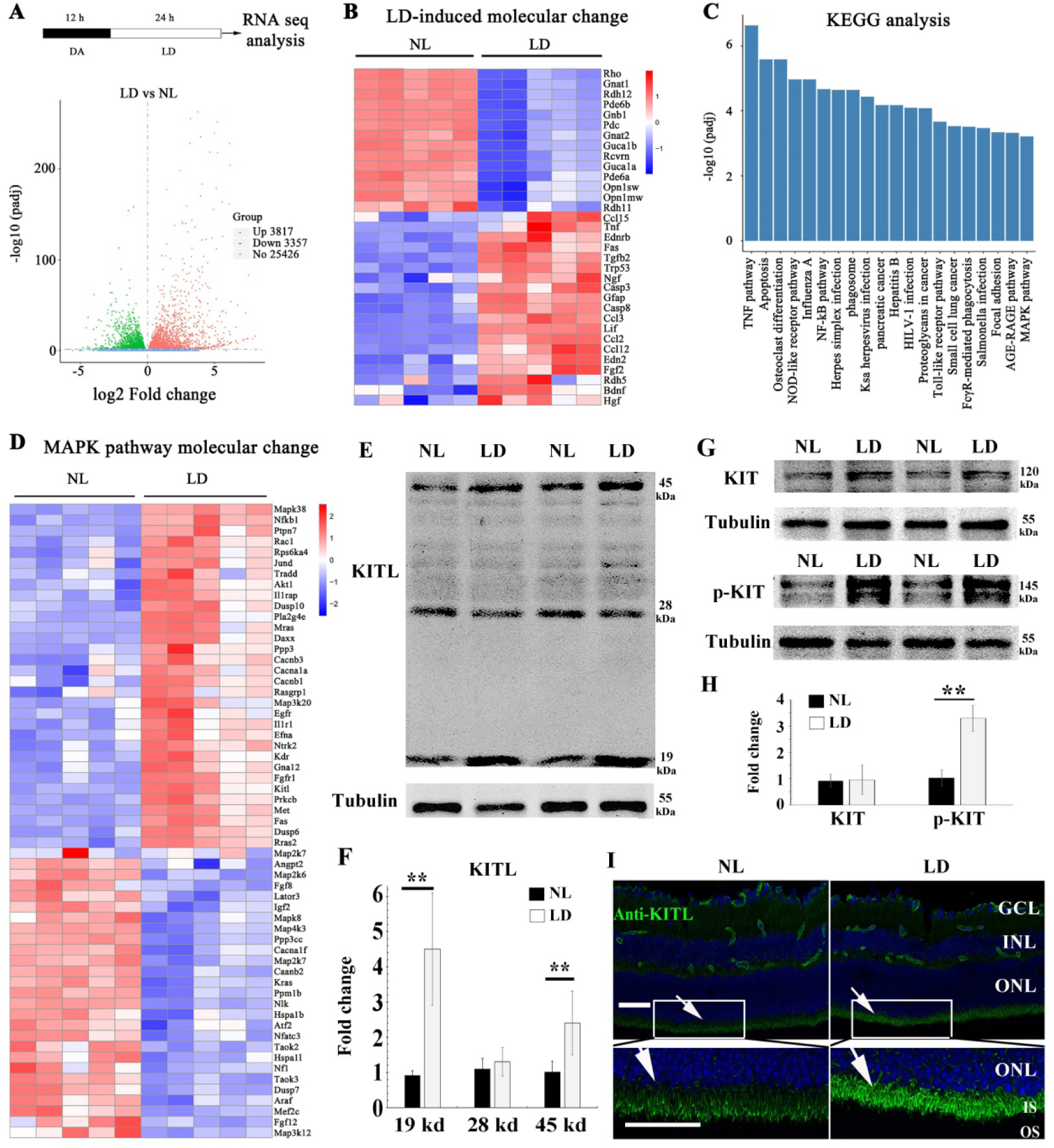
Light damage induces increased expression of KITL and activation of KIT in mouse retina. (**A**) Schematic representation of time frame and analysis of light damage (LD). Two-month-old albino mice were raised in the dark for 12 hours and then exposed to constant white light of 15,000 lux for 1 day to induce retinal damage. The volcano map of transcriptome analysis shows a global view of gene expression. (**B**) Heat map of the selective LD-induced differentially expressed reported genes. The columns for NL (normal light) or LD represent the results from five biological replicates. (**C**) The bar graph of KEGG analysis shows the top 20 differentially signaling pathways. (**D**) Heat map shows the differentially expressed genes of the MAPK pathway. (**E**) Western blotting analysis of KITL in retinas after LD. (**F**) Quantification of western blot bands shows the expression levels of the KITL. Note that both the 19 kDa and 45 kDa isoforms of KITL were upregulated by LD. (**G**) Western blot analysis of the tyrosinase kinase receptor KIT (upper panels) and its phosphorylated form (lower panels) in the retinas after LD. (**H**) Quantification of western blot bands show the expression levels of KIT and its phosphorylation levels. Note that p-KIT was upregulated by LD. (**I**) Immunostaining of light-treated retina detected by anti-KITL antibody. Each image is representative of at least 5 retinas. GCL, ganglion cell layer; INL, inner nuclear layer; IS, photoreceptor inner segments; ONL, outer nuclear layer; OS, photoreceptor outer segments. ** indicates *p*<0.01. Scale bar, 50 μm.

### The *Kit*^*Wps*^ mutation does not cause detectable changes in retinal function and structure but disrupts KIT activation in the retina

To determine whether upregulation of KIT signaling has any functional relevance during excessive light exposure, we used mice harboring a mutant *Kit* allele on a C57BL/6J background, *Kit*^*Wps*^. This allele encodes a KIT protein with an Asp-to-Asn (D-to-N) mutation at residue 60 in the extracellular domain that reduces, but does not eliminate, KIT signaling, which allows *Kit*^*Wps*^ homozygotes to be viable, though they are totally white and infertile (*46*). However, these mice have no detectable changes in the retinal fundus, function and structure, and eye pigmentation is similar to those in *Kit+/+* mice (Fig. 2A). Besides, the amplitudes of a wave and b wave under scotopic or photopic conditions are not significantly different between *Kit+/+* and *Kit*^*Wps/Wps*^ mice (Fig. 2B and C). As expected, these mice do not show structurally abnormal retinas (Fig. 2D). The numbers of cell nuclei in the ganglion cell layer (GCL), inner nuclear layer (INL) and outer nuclear layer (ONL) were similar to those in *Kit+/+* retinas (Fig. 2E). Also, as shown in Fig. 2F, the immunoreactivity of OTX2 (marker for allbipolar cells), PKCα (marker for “On” bipolar cells), Rhodopsin (marker for rod photoreceptor cells) and Opsin (marker for cone photoreceptor cells) was as in *Kit+/+* retinas. Likewise, there was also no significant difference in the number of bipolar cells and cone photoreceptor cells (Fig. 2G). These data show that under normal conditions, the *Kit*^*Wps*^ mutation does not cause detectable changes in retinal function, histology and structure.

**Figure 2.**
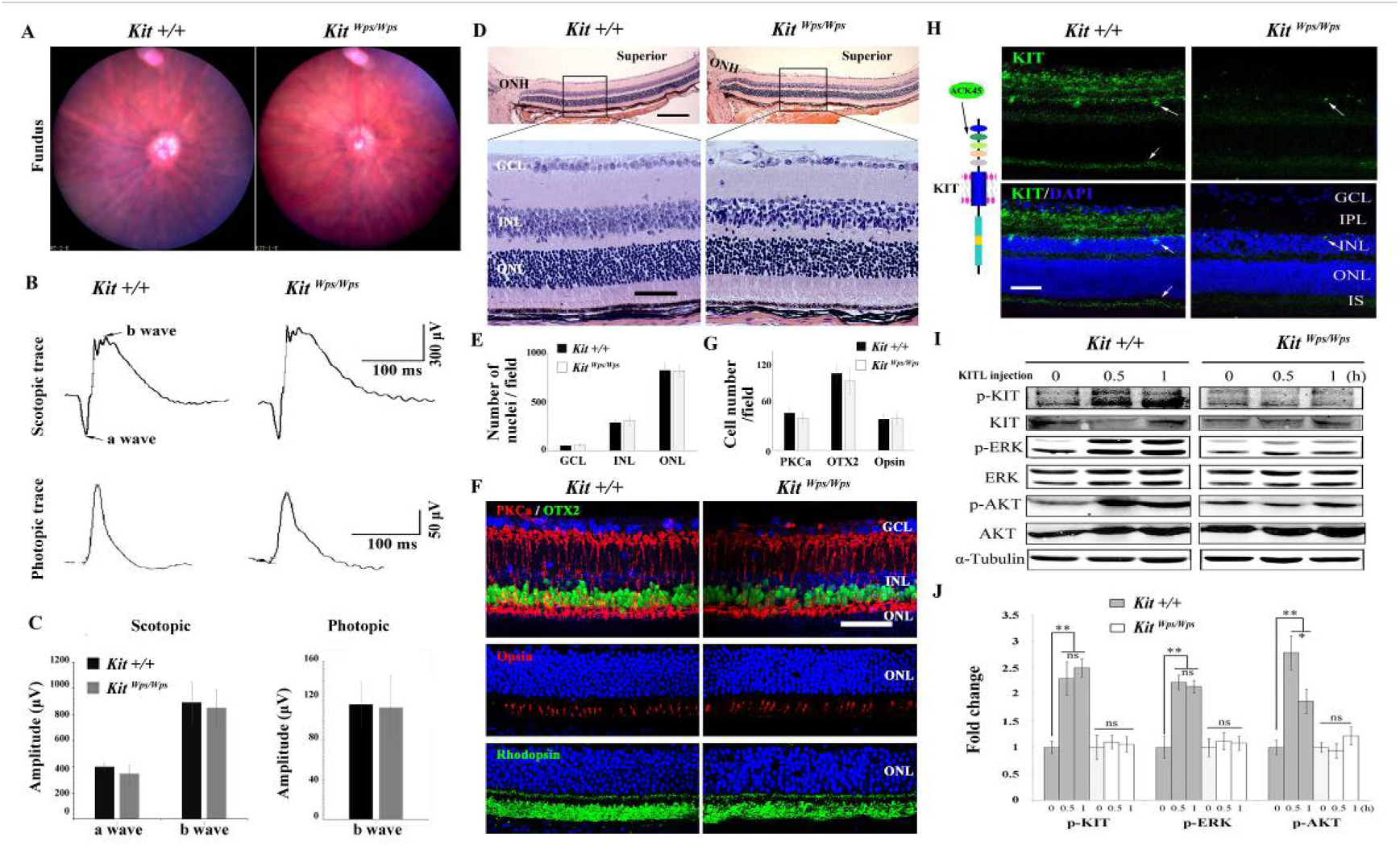
No changes in retinal cell distribution and structure, but disruption of Kit activation in *Kit*^*Wps/Wps*^ retina. (**A**) Fundus images of 2-month-old *Kit+/+* and *Kit*^*Wps/Wps*^ mice. (**B**) ERG traces of *Kit+/+* (left panels) and *Kit*^*Wps/Wps*^ (right panels) retinas under scotopic (upper panels) and photopic conditions (lower panels). (**C**) Quantification of ERG amplitudes under scotopic and photopic conditions. (**D, E**) Histological images of H&E staining from 2-month-old *Kit+/+* and *Kit*^*Wps/Wps*^ retinas (**D**) and quantification showing the mean number of nuclei localized at GCL, INL, and ONL over the length of 0.35 mm between 300 μm to 700 μm from the optic nerve head in dorsal retina of *Kit+/+* and *Kit*^*Wps/Wps*^ mice (n=5) (**E**). Representative histological sections are from superior retinas 0.3 mm from the optic nerve head. (**F**) Immunohistochemistry for OTX2, PKCα, Opsin and Rhodopsin in 2-month-old *Kit+/+* and *Kit*^*Wps/Wps*^ retinas. (**G**) Quantification of OTX2+, PKCα+ and Opsin+ cells in the indicated retinas. Each image is representative for at least 5 retinas. The analyzed sectional numbers are from at least 30 sections from at least 5 retinas. (**H**) Schematic representation of KIT protein structure and the location recognized by the ACK45 antibody (left panels) and immunostaining images of ACK45 antibody in 3-month-old *Kit+/+* and *Kit*^*Wps/Wps*^ retinas (right panels). (**I**) Western blots show phosphorylation of KIT, ERK and AKT in *Kit+/+* and *Kit*^*Wps/Wps*^ retinas after the injection of KITL (5.6 nM) at the indicated time points. (**J**) Quantification of western blot bands shows the phosphorylation levels of KIT, ERK and AKT. GCL, ganglion cell layer; IPL, inner plexiform layer; INL, inner nuclear layer; IS, photoreceptor inner segments; ONL, outer nuclear layer; OS, photoreceptor outer segments. * or ** indicates *p*<0.05 or *p*<0.01. Scale bar, 50 μm.

We next assessed whether the mutation would interfere with KIT signaling in the retina. As shown in Supplemental Figure 2, western blots and immunostaining using the AB5506 antibody (recognizing the intracellular kinase domain of KIT) did not detect differences in KIT expression levels between *Kit+/+* and *Kit*^*Wps*^ homozygous retinas. Nevertheless, immunostaining using the ACK45 antibody (recognizing the extracellular immunoglobulin domain of KIT protein) showed a significantly decreased immunoreactivity in the photoreceptor layer, the inner plexiform layer and the ganglion cell layer (Fig. 2H). Because the point mutation lies in the first immunoglobulin domain of KIT, which is required for the interaction with KITL, we hypothesized that it would impair KIT activation by KITL. Indeed, upon intravitreal injection of KITL, phosphorylation of KIT and its downstream targets ERK and AKT were prominent in *Kit+/+* but not in *Kit*^*Wps/Wps*^ retinas (Fig. 2I and J). These results indicate that the mutation substantially reduced activation of KIT by KITL in the retina.

### Disruption of KIT signaling aggravates retinal dysfunction and exacerbates retinal degeneration after prolonged light exposure

To gain insights into the role of KIT signaling specifically during LD of the retina, we used mice homozygous for *Kit*^*Wps*^ on C57BL/6J background. Even though C57BL/6J mice express the Leu450-Met variant of RPE65 and are resistant to LD (*48*), they have been used as a suitable model of light-induce retinal damage (LIRD) because their susceptibility to LD is similar to that of humans (*49*).Thus, we first dark-adapted *Kit+/+* and *Kit*^*Wps/Wps*^ mice and examined them by ERG. Subsequently, the mice were light-exposed (15,000 lux) for 8 days, with the dilated pupils by a mydriatic agent once a day, and then re-examined by ERG (Fig. 3A). As shown in Fig. 3B to D, under scotopic conditions (rod response), the a- and b-wave amplitudes were similar in *Kit+/+* and *Kit*^*Wps/Wps*^ mice under normal light (426 ± 58μV and 837 ± 1314μV, respectively, in *Kit+/+*, n=8; and 439 ± 83μV and 863 ± 114μV, respectively, in *Kit*^*Wps/Wps*^ mice, n=8). After LD, the a- and b-wave amplitudes declined in *Kit+/+* to 294 ± 37μV and 521 ± 27μV, respectively (n=8) and in *Kit*^*Wps/Wps*^ mice to 188 ± 80μV and 365 ± 63μV, respectively (n=8). These data indicate that the rod-driven circuit was damaged more severely in *Kit*^*Wps/Wps*^ retinas compared to *Kit+/+* retinas. Similarly, under photopic conditions (cone response) (Fig. 3, E and F), the b-wave ERG responses were weakened in *Kit*^*Wps/Wps*^ mice more severely (from 89.4 ± 15μV to 40 ± 10μV, n=8) than those in *Kit+/+* mice (from 95 ± 13μV to 61 ± 9μV, n=8), indicating that the cone-driven circuit was also damaged more severely in *Kit*^*Wps/Wps*^ retinas. These results suggest that normal KIT signaling helps to protect against retinal damage following LD.

**Figure 3.**
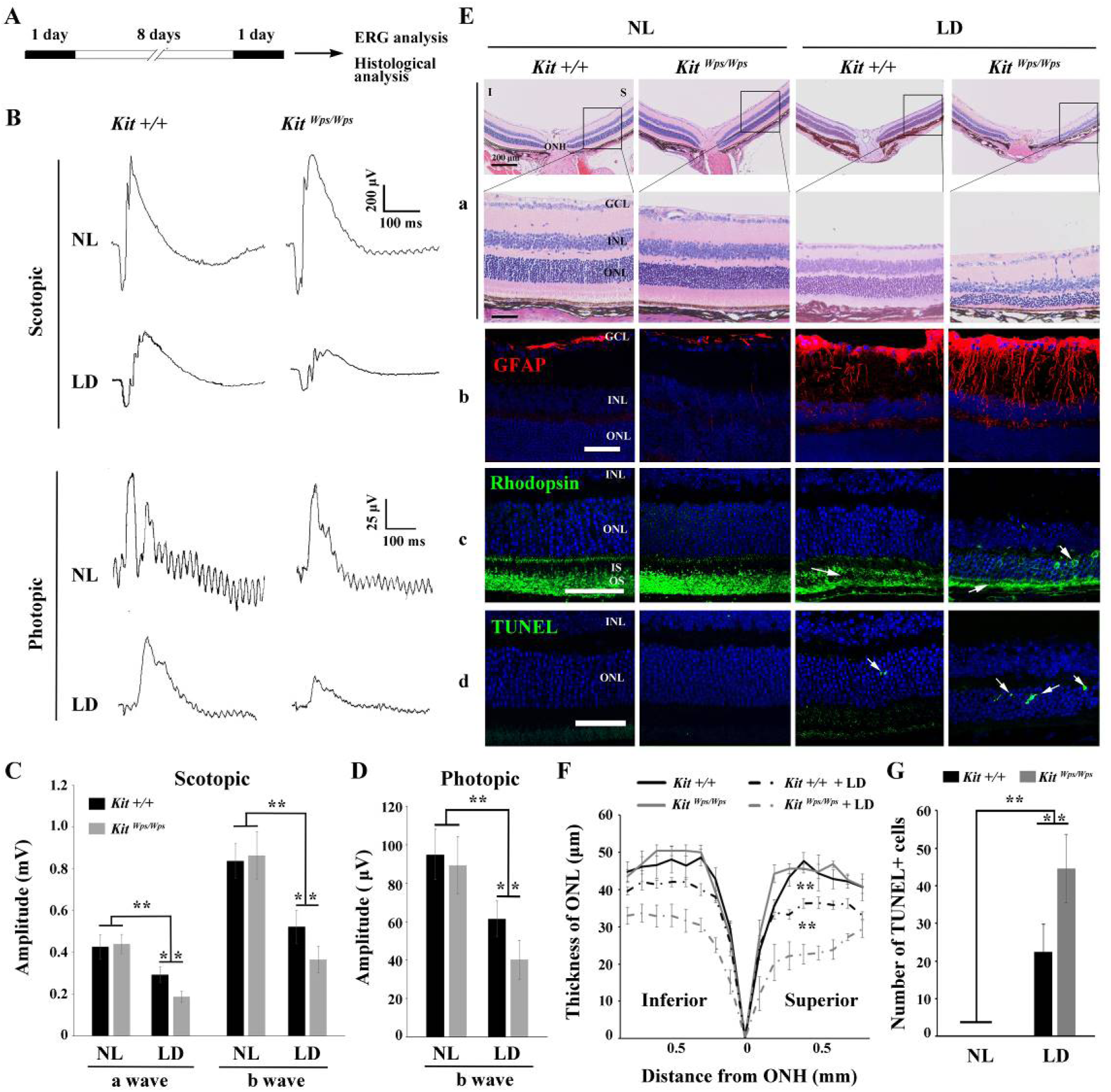
*Kit*^*Wps*^ mutation significantly damages retinal function and accelerates retinal degeneration during LD. (**A**) Schematic representation of LD treatment of mice for D-F. Three-month-old *Kit+/+* and *Kit*^*Wps/Wps*^ mice were raised in the dark for one day, and then exposed to constant high intensity light of 15,000 lux for 8 days with mydriasis treatment to induce retinal damage. Retinal functions were subsequently assessed by electroretinography (ERG) after a one-day dark adaptation. (**B**) ERG traces of *Kit+/+* (left panels) and *Kit*^*Wps/Wps*^ (right panels) retinas before and after LD treatment under scotopic (upper panels) and photopic conditions (lower panels). Note that both scotopic and photopic responses of the *Kit*^*Wps/Wps*^ retinas were similar to those of *Kit+/+* retinas with normal light (NL). Usually, the scotopic ERG result is shown as step-wise figure. (**C** and **D**) Quantification of ERG amplitudes under scotopic **(C)** and photopic conditions (**D**). Note that both scotopic and photopic responses of *Kit*^*Wps/Wps*^ retinas were impaired more severely than those in *Kit+/+* retinas. Each trace is the average of individual records from at least 5 mice. (**E**) Retinal degeneration analysis of *Kit+/+* and *Kit*^*Wps/Wps*^ mice before and after LD treatment by HE staining (**a**), GFAP staining (**b**), Rhodopsin staining (**c**) and TUNEL detection (**d**). Arrows point to abnormal translocation signal of the rhodopsin (**c**) and dead cells (**d**). (**F**) The curve diagram shows the thickness of ONL from *Kit+/+* and *Kit*^*Wps/Wps*^ retinas under NL or high intensity LD conditions. (**G**) Quantification of the number of TUNEL positive cells in the ONL. GCL, ganglion cell layer; I, inferior; INL, inner nuclear layer; IS, photoreceptor inner segments; ONL, outer nuclear layer; S, superior. ** indicates *p*<0.01. Scale bar, 50μm.

Retinal degeneration analysis revealed that along with the reduction of the electrical responses, LD also led to significant morphological changes, reactive gliosis, Rhodopsin translocation, and photoreceptor cell death. As shown in Fig. 3E, compared to light-damaged *Kit+/+* retinas, similarly treated *Kit*^*Wps/Wps*^ retinas showed a severely reduced thickness of the ONL both in superior and inferior regions. For reactive gliosis analysis, in unstressed *Kit+/+* or mutant retinas, GFAP immunoreactivity was only detected in the ganglion cell layer, while Müller glial cells became positive for GFAP, but much more so in *Kit*^*Wps/Wps*^ compared to *Kit+/+* mice after light stress (Fig 3E, panels b). Likewise, rhodopsin immunoreactivity was not only more severely reduced in light-damaged *Kit*^*Wps/Wps*^ compared to *Kit+/+* retinas but also showed abnormal presence in the ONL (Fig. 3E, arrows in panels c). These results were consistent with TUNEL assays to evaluate cell death. As shown in Fig. 3E, panels d, although dead cells were present in the ONL of *Kit+/+* retinas (22 ± 7 per section, n=10), their number was much higher in *Kit*^*Wps/Wps*^ retinas (44 ± 9, n=10). Hence, these data show that after excessive light exposure, the photoreceptor cells of *Kit*^*Wps/Wps*^ mice are not only structurally but also functionally impaired to a greater extent than those of *Kit+/+* mice. This finding indicates that the *Kit*^*Wps*^ mutation accelerates photoreceptor cell degeneration during LD.

### KITL prevents photoreceptor cell death-associated with LD

As the above results indicated a role for KIT signaling in light-induced photoreceptor cell degeneration, we asked whether KITL may plays a protective role in LD. Hence, we overexpressed KITL by intraocular injection of an engineered AAV virus. We first tested in which retinal areas an AAV8-GFP vector is expressed by scanning fluorescent ophthalmoscopy and by optical coherence tomography. After virus infection for 14 days, fundus photographs showed that AAV-GFP infected nearly the entire retina, and cross section showed strong GFP signals in photoreceptor cells (Supplemental Figure 3).We then ectopically expressed KITL using an appropriately engineered viral expression vector, AAV8-KITL (schematically shown in Fig. 4A). Two weeks after injection, western blots showed significant high expression of KITL (18.9 ± 3.5 folds, n=4) that by immunostaining was almost entirely restricted to photoreceptors (Fig. 4B and C), suggesting that overexpression of KITL can be achieved successfully in photoreceptor cells.

**Figure 4.**
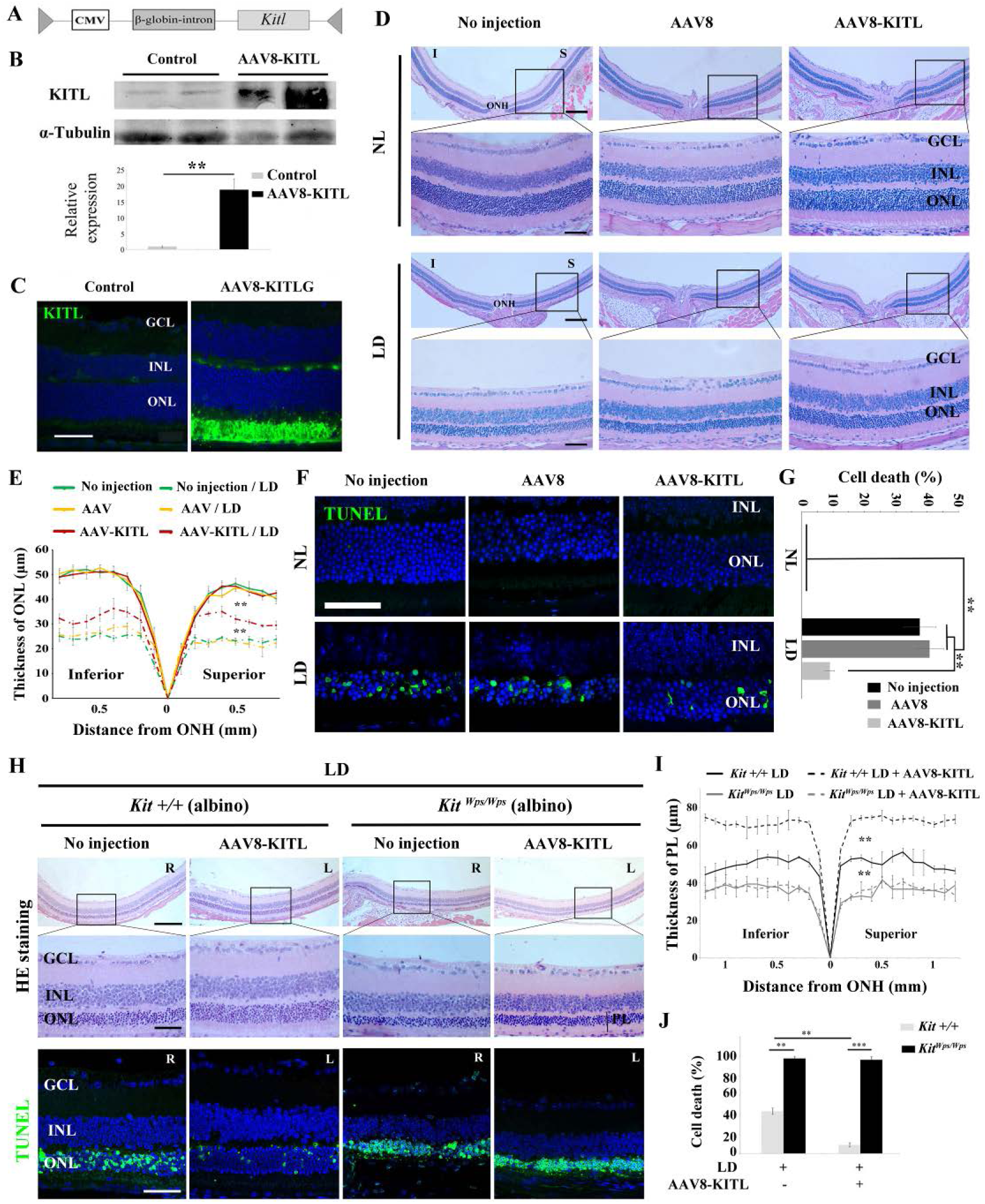
Overexpression of KITL by AAV8-KITL virus prevents light-induced retinal degeneration. (**A**) Schematic representation of the AAV8-KITL construct. (**B**) Western blots show the expression of KITL in the retina 2 weeks after intraocular injection of AAV8-KITL. The bar graph shows the relative expression of KITL based on the results of western blots. (**C**) Immunostaining images show the immunoreactivity of KITL in 3-month-old *Kit+/+* retinas 2 weeks after intraocular injection of AAV8-KITL. (**D**) Histological images of H&E staining from 3-month-old albino retinas infected with AAV8 or AAV8-KITL virus and kept under NL (upper panels) or high intensity LD (15,000 lux) continuously for 3 days (lower panels). (**E**) The curve diagram shows the thickness of ONL from albino retinas under NL or high intensity LD conditions. (**F, G**) Images of TUNEL assays from the retinas (**F**) and bar graph show the cell death rate of photoreceptor cells (**G**) under the indicated conditions. (**H**) Retinal degeneration analysis of 3-month-old *Kit*^*Wps/Wps*^; albino retinas infected with or without AAV8-KITL virus and kept under high intensity LD by HE staining (upper panels) and TUNEL detection (down panels). (**I**) The curve diagram shows the thickness of the photoreceptor cell layer from the indicated retinas. (**J**) Quantification of the rate of cell death in the ONL of the indicated retinas. GCL, ganglion cell layer; INL, inner nuclear layer; ONL outer nuclear layer; ONH, optic nerve head; PL, photoreceptor cell layer. * or ** indicates *p*<0.05 or *p*<0.01. Scale bar: 50 μm.

Using albino mice, we then tested whether overexpression of *Kitl* would affect retinal damage after light exposure. Without LD, neither AAV8 (Supplemental Figure 4A) nor AAV8-KITL caused detectable changes of retinal morphology (Fig. 4 D and E). After 3 days of LD treatment, retinas infected by AAV8-KITL virus were thicker than those infected by AAV8 virus or left uninjected (Fig. 4D and E). Furthermore, TUNEL analysis showed that injection of AAV8 virus or AAV8-KITL virus did not cause photoreceptor cell death under normal light conditions, while photoreceptor cell death was less pronounced in retinas infected by AAV8-KITL virus (9 ± 1.6%, n=6) than those infected by AAV8 virus (40.8 ± 4.5%, n=6) or left uninjected (37.7 ± 5%, n=6) under LD (Fig. 4F and G). These results suggest that AAV8-KITL virus prevents light-induced photoreceptor cell death. To confirm that the protective role of AAV8-KITL virus is mediated by overexpression of KITL, we examined KITL expression in the retinas after LD treatment. Indeed, the data of immunostaining showed high expression levels of KITL in photoreceptor cells infected by the AAV8-KITL virus compared with uninjected retinas (Supplemental Figure 4B). Taken together, these data indicate that overexpression of KITL prevents light-induced photoreceptor cell degeneration.

To investigate whether protection of AAV8-KITL virus indeed depends on KIT, we used *Kit*^*Wps/Wps*^ albino mice through crossing between *Kit*^*Wps*^ heterozygotes and BALB/c albino (*Tyr*^*c*^/*Tyr*^*c*^) mice. After 3 days of LD treatment, the retinas in both *Kit+/+* albino and *Kit*^*Wps/Wps*^ albino mice showed reduced thickness and near-absence of outer segments, but the effects were more severe in *Kit*^*Wps/Wps*^ albino compared to *Kit+/+* albino mice (Fig. 5, H and I). Importantly, injection of the AAV8-KITL virus prevented light-induced reduction of retinal thickness in *Kit+/+* albino mice, but, as expected, not in *Kit*^*Wps/Wps*^ albino mice (Fig. 5, H and I). Consistent with these results, TUNEL assay showed a cell death rate of 43.6 ± 3.3% in *Kit+/+* albino mice and 97.9 ± 1.9% in *Kit*^*Wps/Wps*^ albino mice (n=4) (Fig. 5H and J), while infection of AAV8-KITL significantly reduced the death rate of photoreceptor cells to 8.9 ± 2.2% in *Kit+/+* albino mice but not in *Kit*^*Wps/Wps*^ albino mice, where it remained high (96.7 ± 3.2%, n=4). Collectively, these data suggest that overexpression of KITL can prevent photoreceptor cell death from LD through activation of the receptor KIT.

**Figure 5.**
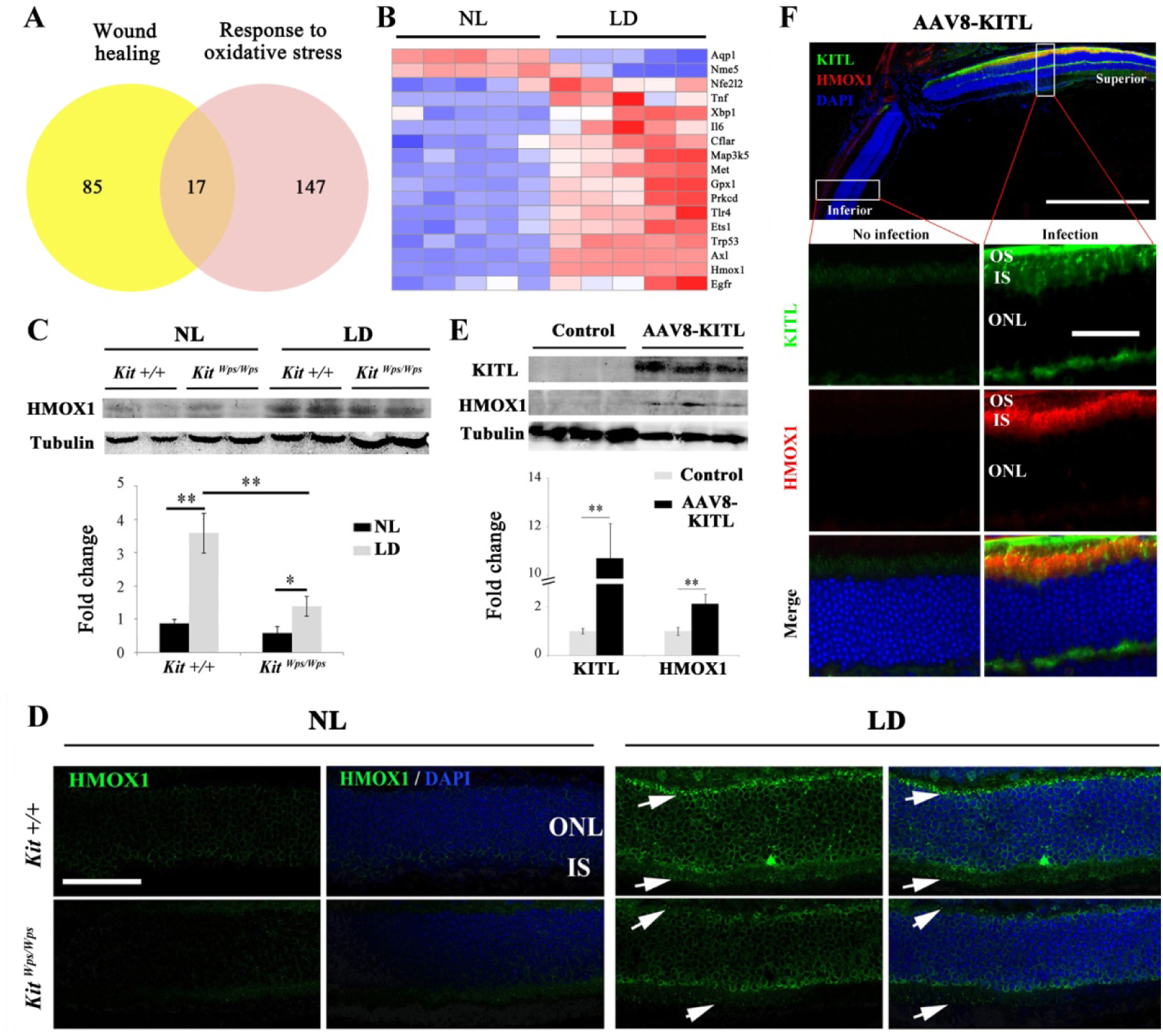
KIT signaling regulates expression of *Hmox1* in photoreceptor cells. (**A**) Venn diagram shows numbers of differentially expressed genes response to oxidative stress and wound healing. (**B**) Heat-map of differentially expressed genes from the 17 genes common between the wound healing group and response to oxidative stress group. Messenger RNA expression changes of anti-oxidant genes in *Kit+/+* retinas after LD for 1 day. (**C** and **D**) Expression of HMOX1 in *Kit+/+* and *Kit*^*Wps/Wps*^ retinas were detected by western blots and immunostaining. Note that the expression of HMOX1 was more significantly upregulated in light-degenerated *Kit+/+* compared to light-degenerated *Kit*^*Wps/Wps*^ photoreceptors. (**E**) Western blots show the expression of KITL and HMOX1 in the retinas infected with control AAV8 or AAV8-KITL virus. Bar graphs show the relative expression of KITL and HMOX1 in retinas based on the results of western blots. (**F**) Immunostaining images show immunoreactivity of KITL and HMOX1 in the retina one week after subretinal injection of AAV8-KITL. IS, photoreceptor inner segments; ONL, outer nuclear layer; OS, photoreceptor outer segment. * or ** indicates *p*<0.05 or *p*<0.01. Scale bar: 50 μm.

### KIT signaling regulates *Hmox1* expression in photoreceptor cells

Studies of human disease and animal models of photoreceptor degeneration have shown that photoreceptor loss is often accompanied by oxidative damage, including irreversible oxidation of DNA and other biomolecules (*50*). Hence, in order to analyze the mechanism by which LD leads to the demise of photoreceptor cells, we focused on oxidative damage. Hence, we analyzed whether a series of antioxidant and wound healing genes would show expression changes in *Kit+/+* retinas following LD (Fig. 5A). In fact, after LD, the expression of 17 such genes was changed in *Kit+/+* retinas including *Axl* and *Hmox1* (Fig. 5B). The protein product of *Hmox1*, HMOX1, catalyzes the degradation of heme into three biologically active end products, biliverdin/bilirubin, CO and ferrous ion (*51*). It is known to be induced by oxidative stress in multiple tissues as well as by LD of the retina (*52*) and thought to act as an antioxidant (*51*).Under normal light/dark cycle conditions, HMOX1 levels analyzed on western blots (Fig. 5C) or by immunostaining (Fig. 5D) were similarly low in both *Kit+/+* and *Kit*^*Wps/Wps*^ retinas. After LD, however, HMOX1 expression was significantly upregulated in *Kit+/+* retinas (3.6±0.6 fold) but only slightly in *Kit*^*Wps/Wps*^ retinas (1.4± 0.5 fold, n=3) (Fig. 5C and D). These data indicate that LD-dependent upregulation of *Hmox1* is markedly influenced by KIT signaling.

The importance of KIT signaling for the regulation of HMOX1 expression can also be demonstrated *in vitro* in 661W photoreceptor cells. To test for the differential effects of *Kit+/+* and mutant KIT, we transfected such cells with a control vector or vectors encoding Kit-WT or Kit-Wps cDNA and treated the cells with KITL. As expected, the ACK2 antibody, which reacts with the cytoplasmic portion of KIT, readily detects KIT after transfection of either of the two KIT vectors (Supplemental Figure 5A). Western blots indicated an increase of 2.9 ± 0.23 fold with Kit-WT and 2.2 ± 0.9 fold with Kit-Wps (Supplemental Figure 5B).Transfection with KIT also led to upregulation of HMOX1, although with a clear difference between KIT-WT (6.2 ± 0.86 fold, n=4) and Kit-Wps (3 ± 0.6 fold, n=4). These results are consistent with the *in vivo* results described above.

Given the above *in vitro* results, we then asked whether ectopic activation of KIT signaling would induce *Hmox1* expression in the retina without LD. To answer this question, we analyzed the expression of HMOX1 in retinas of *Kit+/+* mice kept under normal light conditions and infected with AAV8-KITL virus. One week after injection, western blots showed a correlation between high levels of KITL (10.7 ±1.4 fold, n=4) and HMOX1 (2.1 ± 0.4 fold, n=4) (Fig. 5E). Consistent with this observation, immunostaining showed that the expression of KITL was dramatically enhanced in the superior retinal region around the location of injection, coinciding with high level HMOX1 expression (Fig. 5F). These data suggest that activation of KIT signaling can induce HMOX1 expression in *Kit+/+* retinas independent of LD treatment.

### KITL/KIT signaling activates NRF2 to regulate *Hmox1* expression

To understand how KIT regulates *Hmox1* expression, we analyzed whether KIT signaling is able to regulate the expression of transcript factor NRF2, an upstream regulator of *Hmox1* that is activated in many cell types in response to stressful environments (*53*). Although western blots showed no change in *Nrf2* expression in retinas after light treatment (Fig. 6A and B), immunostaining indicated a dramatic increase of NRF2 in the photoreceptor nuclei after light treatment for 8 days (Fig. 6C). Intriguingly, this nuclear accumulation was much reduced in *Kit*^*Wps/Wps*^ mice (14 ± 5 per section, n=6) compared to *Kit+/+* mice (28 ± 9, n=6) (Fig. 6D). These results suggest that Kit signaling regulates nuclear translocation of NRF2 in photoreceptor cells.

**Figure 6.**
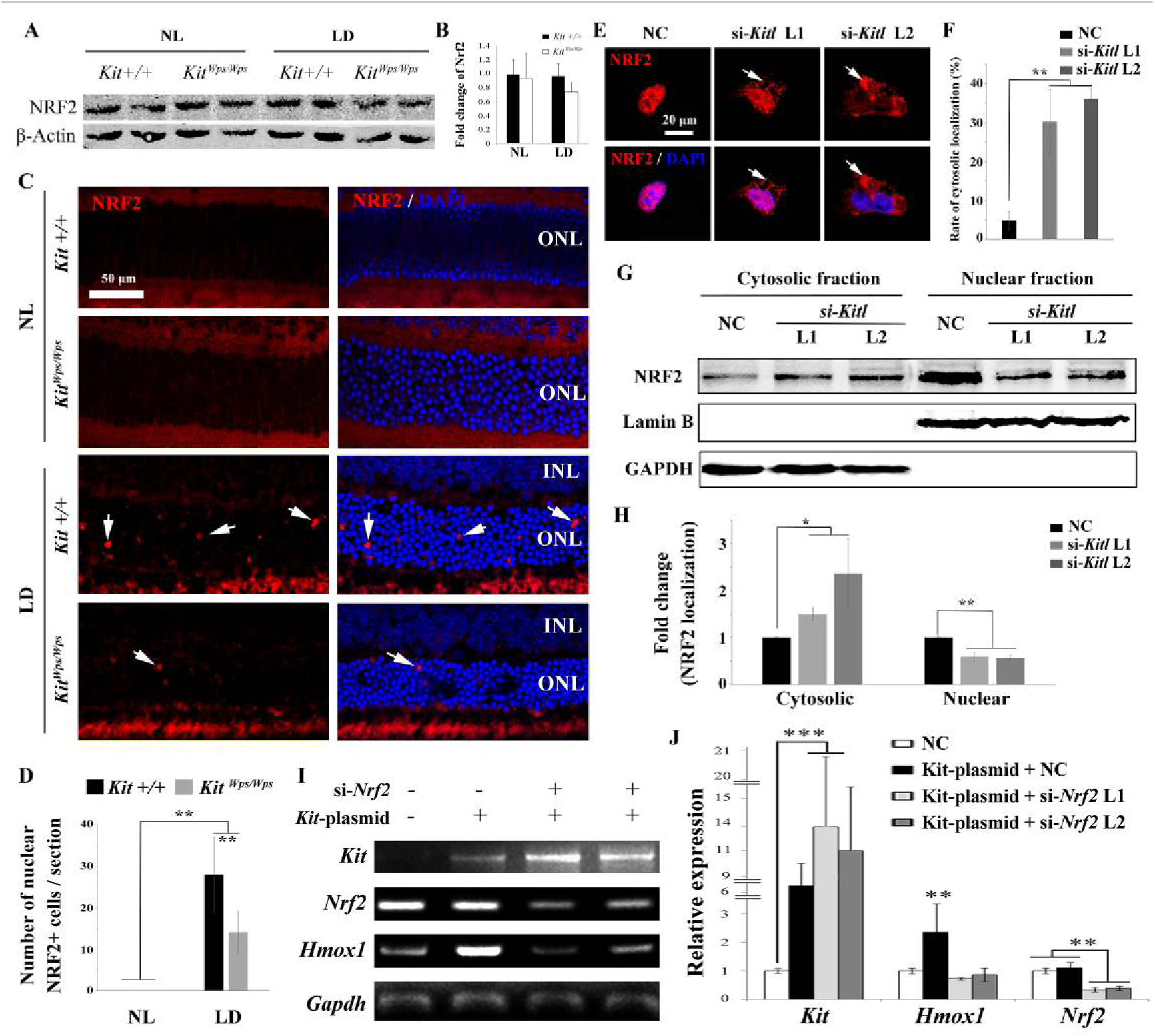
KIT signaling acts through the transcription factor NRF2 to regulate *Hmox1* expression. **(A**) Western blots for analyzing NRF2 expression in *Kit+/+* and *Kit*^*Wps/Wps*^ retinas under the indicated conditions. (**B**) The bar graph shows quantification of NRF2 expression in the indicated retinas. (**C**) Immunostaining of anti-NRF2 in *Kit+/+* and *Kit*^*Wps/Wps*^ retinas under the indicated conditions. The arrows point to nuclear signals of NRF2 in the ONL. (**D**) Quantification of the number of nuclear NRF2 positive cells in the retina. Note that LD induced nuclear accumulation of NRF2 in photoreceptor cells. (**E-H**) Analysis of subcellular localization of NRF2 in 661W photoreceptor cells treated with si-*Kitl* by immunostaining (**E**, and **F**) and western blot (**G**, and **H**). Note that knockdown of *Kitl* led to an increase in the proportion of cells with cytosolic NRF2. (**I, J**) Analyses of the regulation of HMOX1 in 661W cells after overexpression of KIT together with si-*Nrf2*. The images of RT-PCR (**I**) and qPCR (**J**) show the expression levels of *Kit*, *Nrf2*, and *Hmox1* under the indicated treatments. Note that upregulation of *Hmox1* induced by overexpression of KIT was blocked by the knockdown of *Nrf2*. * indicates *p*<0.05, ** indicates *p*<0.01. NS indicates no significant difference.

To further examine whether NRF2 nuclear accumulation was dependent on KITL, we used Kitl-specific siRNAs to reduce expression of *Kitl* in 661W photoreceptor cells (Supplemental Figure 6). Indeed, immunostaining revealed that the both Kitl-si-RNAs increase the number of cells with cytosolic NRF2 expression from 4.74 ± 2.27% to 30.2 ± 8.3% or 36.8 ± 2.24 % (n=3), respectively (Fig. 6 E and F). In addition, western blots showed that knockdown of *Kitl* increased the levels of NRF2 in the cytoplasm (1.5 ± 0.13 fold, n=3) and decreased the levels of nuclear NRF (20.59 ± 0.09 fold, n=3) (Fig. 6G and H). Taken together, these data clearly show that interference of KIT signaling disturbs the nuclear accumulation of NRF2 in photoreceptor cells.

To investigate whether the regulation of *Hmox1* by KIT signaling depends on NRF2, we then used siRNA to knockdown *Nrf2* in 661W cells overexpressing KIT (Fig. 6I and J). Consistently, overexpression of KIT did not significantly change *Nrf2* expression (1.1 ± 0.2 fold, n=3) but upregulated *Hmox1* expression (2.35 ± 1 fold, n=3). This upregulation was blocked by knockdown of *Nrf2*. Collectively; these data suggest that KIT signaling activates NRF2 to regulate the expression of *Hmox1* in photoreceptor cells.

### *Hmox1* overexpression partially rescues retinal degeneration in light-damaged *Kit* mutant mice

While the above results demonstrated that KIT signaling regulates *Hmox1* expression in the retina and plays a neuroprotective role during LD, the relevant function of HMOX1 in retina *in vivo* was still unclear. To analyze whether KIT signaling exerts its protective effect against LD through HMOX1, we ectopically expressed *Hmox1* in *Kit*^*Wps/Wps*^ albino mice, using an AAV8-based vector engineered to express HMOX1 (Fig. 7A). Western blotting and immunostaining showed photoreceptor-restricted expression after 2 weeks (Fig. 7A and B), with levels similar to those observed in uninjected *Kit+/+* retinas after LD. ERG measurements showed that without LD, control AAV8 or AAV8-Hmox1 virus did not cause significant changes in the ERG scotopic trace (Fig. 7C). Following 3 days of LD, however, the amplitudes of ERG scotopic traces were depressed after infection with AAV8 or AAV8-HMOX1 virus, but the b-wave amplitudes were higher when AAV8-HMOX1 was used (381 ± 40μV, n=5) compared to when control AAV8 was used (152 ± 33 μV, n=5) (Fig.7D). These results suggest that overexpression of HMOX1 partially rescues retinal function in the *Kit*^*Wps/Wps*^ albino mice during LD. Furthermore, histological analyses showed that after LD, AAV8-HMOX1-infected retinas were thicker than control AAV8-infected ones, especially with respect to the photoreceptor cell layer (Fig. 7E). These data suggest that overexpression of HMOX1 protects photoreceptor cells in *Kit*^*Wps/Wps*^ albino retinas against LD and that LD-induced HMOX1 plays an important antioxidant role in retinas in vivo.

**Figure 7.**
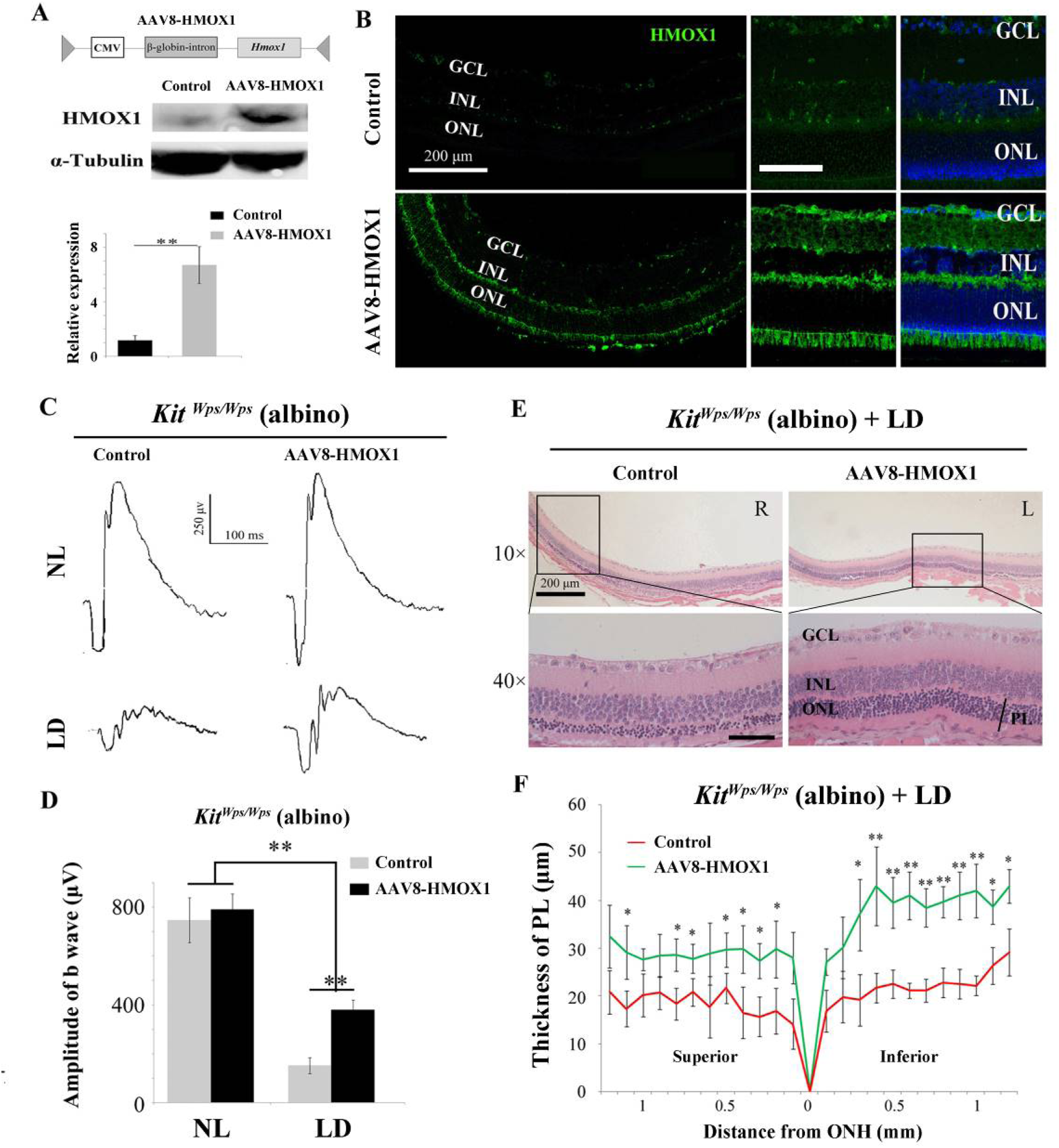
Protective role of HMOX1 in LD-treated *Kit*^*Wps/Wps*^ retina. (**A**) Schematic representation of the AAV8-HMOX1 construct (upper panel) and western blots show the expression of KITL in the retina 2 weeks after subretinal injection of AAV8-HMOX1 virus (lower panels). The bar graph shows the relative expression of HMOX1 based on the results of western blots. (**B**) Ectopic HMOX1 expression in retina by infection of AAV8 or AAV8-HMOX1 virus was visualized by immunofluorescence (n=4). (**C**) ERG scotopic traces of *Kit*^*Wps/Wps*^; albino mice infected with control AAV8 (n=5) or AAV8-HMOX1 (n=5) virus and kept under NL or high intensity LD (15,000 lux) for 3 days. (**D**) Quantification of amplitude of b-wave from standard response based on the results from C. (**E**) Histological analysis of *Kit*^*Wps/Wps*^; albino retina infected with control AAV8 (n=6) or AAV8-HMOX1 (n=6) virus after LD. (**F**) Curve diagram showing the total thickness of the ONL under the indicated conditions. GCL, ganglion cell layer; INL, inner nuclear layer; ONL, outer nuclear layer; PL, photoreceptor cell layer. * or ** indicates *p*<0.05 or *p*<0.01. Scale bar: 50 μm.

To understand whether protection of KITL depends on HMOX1, we used si-*Hmox1* to knockdown the expression of *Hmox1* in 661W photoreceptor cells under light stress. Western blots showed that both Hmox1-si-RNAs downregulate *Hmox1* expression by 0.5 ± 0.2 fold and 0.46 ± 0.23 fold (n=3), respectively (Supplemental Figure 7A and B). We then investigated whether knockdown of *Hmox1* affects light-induced photoreceptor cell death. As shown in Supplemental Figure 7C and D, cell numbers were similar in each group before LD but dramatically decreased in control group (from 409 ± 44 to 147 ± 15 (n=3) after 8 hours of LD treatment). Addition of exogenous KITL (5.6 nM) was able to prevent cell loss caused by LD to 278 ± 28 (n=3). This preventive effect of KITL was totally blocked by knockdown of *Hmox1* via Hmox1-si-RNAs (cell loss 147 ± 16, n=3 and 131 ± 17, n=3 respectively). In addition, the data of propidium iodide (PI) staining also showed that stimulation of KITL decreased light-induced cell death rate from 83 ± 4.6% to 40 ± 5.3% (n=3), but this protection by KITL disappeared when the expression of *Hmox1* was knocked down by Hmox1-si-RNAs (Supplemental Figure 7 C and E). These results suggest that protection of KITL against light-induced photoreceptor cell loss depends on HMOX1.

### KITL preserves photoreceptors and restores retinal function in genetic mouse models of retinal degeneration

We then evaluated whether KITL might provide neuroprotection in mouse models of human RP. Mice homozygous for a mutation inthe gene encoding the photoreceptor cGMP phosphodiesterase, *Pde6b*^*rd10*^ (hereafter called*rd10/rd10)* share many characteristics of human retinal degeneration resulting from a mutation in the homologous gene (*54*). Due to the fact that retinas of *rd10/rd10* mice degenerate soon after birth, we performed subretinal injection of control AAV8 or AAV8-KITL virus into the eyes of such mice at postnatal day 3 and examined their retinal functions at postnatal day 24. As shown in Fig. 8A, ectopic expression of KITL was observed at 3 weeks in *rd10/rd10* photoreceptor cells in the eye infected with AAV8-KITL, but not in those of the control AAV8-infected contralateral eye. These results suggest that subretinal injection of AAV8-KITL virus at P3 can mediate overexpression of KITL in photoreceptor cells of *rd10/rd10* mice. We then analyze the effects of KITL overexpression on retinal degeneration. The thickness of the ONL was significantly increased from the peripheral region to the central region after infection withtheAAV8-KITL virus (Fig. 8B and C). In P24 *rd10/rd10* retinas, Rhodopsin was abnormally translocated from the OS to the ONL, a phenomenon that was partially rescued by infection withtheAAV8-KITL virus (Fig. 8D). TUNEL analysis showed a large number of dead cells in the ONL of *rd10/rd10* mice infected with control AAV8 virus (death rate, 16.2±2%, n=5), but a significantly reduced number after AAV8-KITL virus infection (8.9± 1.6%, n=5) (Fig. 8E). We next examined the effects of KITL on retinal function of *rd10/rd10* mice. The ERG analysis revealed that the scotopic traces of control eyes presented nearly horizontal lines (b-wave amplitude, 162.2±64.4μV, n=5), while infection with the AAV8-KITLvirus led to clear a- and b-waves (b-wave amplitude, 384±146.4μV, n=5) (Fig. 8, F and G), suggesting that overexpression of KITL partially restores the retinal function in *rd10/rd10* mice.

**Figure 8.**
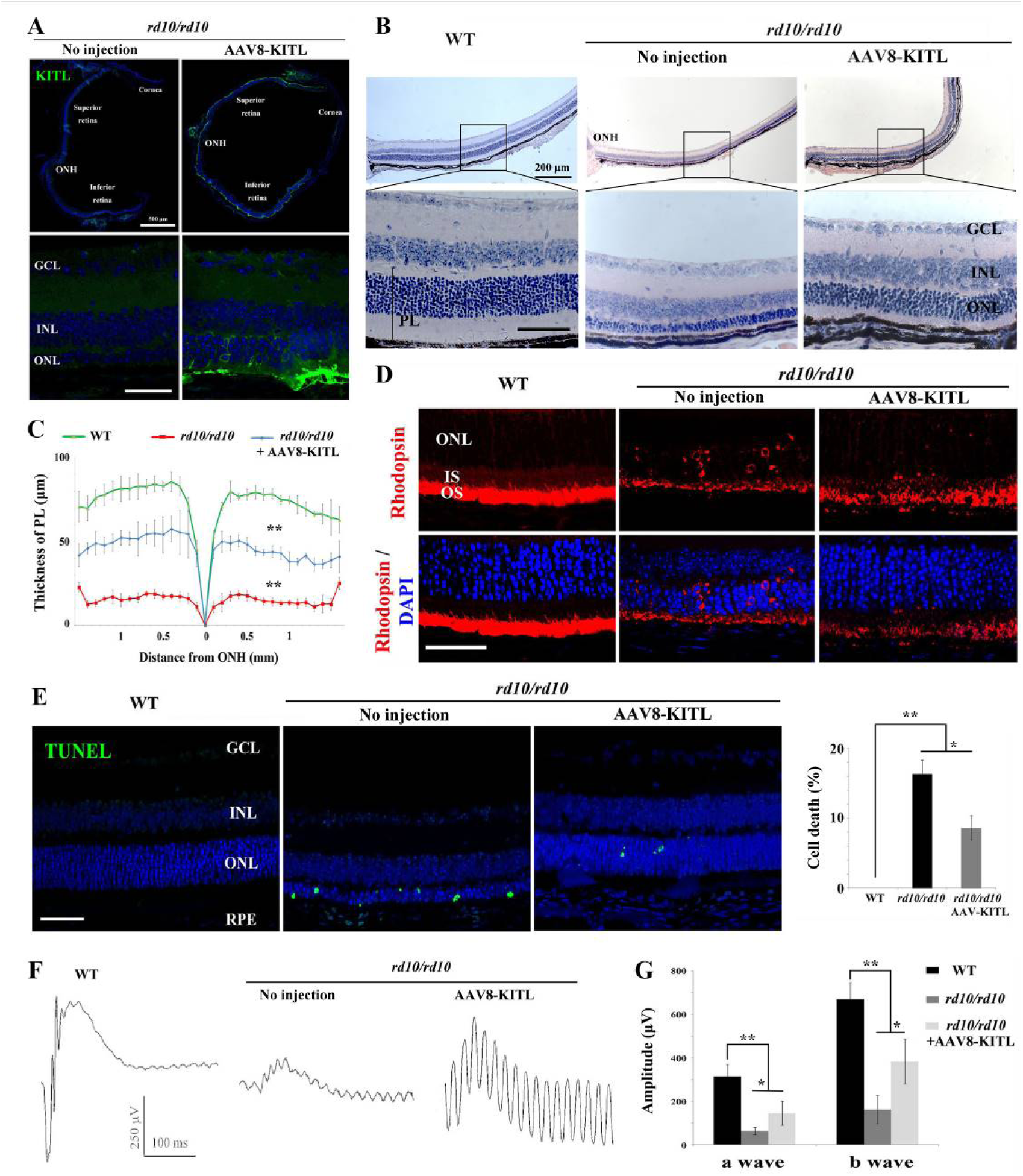
Ectopic expression of KITL prevents retinal degeneration of *rd10* homozygous mice. (**A**) Expression of KITL in *rd10/rd10* retinas 2 weeks after infection with AAV8-KITL virus or control was visualized by immunofluorescence. Note that the subretinal injection was performed in *rd10/rd10* mice at postnatal day 3. (**B** and **C**) Histological analysis of *rd10/rd10* retinas infected with AAV8-KITL virus (n=5) or control (**B**). Curve diagram showing the thickness of total photoreceptor cell layer from *rd10/rd10* retina under the indicated conditions (**C**). **(D)** Rhodopsin expression in WT and *rd10/rd10* retina under indicated conditions. (**E**) Apoptosis of photoreceptor cells from *rd10/rd10* mice with of AAV8-KITL or control was analyzed by TUNEL assay and presented as fluorescent images (upper panels) and as bar graphs after quantitation (lower panels). (**F, G**) ERG scotopic traces (**F**) obtained from *rd10/rd10* mice with infection of AAV8-KITL virus (n=5) or control (n=5) and kept under normal condition for 25 days (right panels). Bar graph (**G**) shows the quantification of the amplitude of the b-wave from standard response based on the results of the ERG scotopic traces (**F**). GCL, ganglion cell layer; INL, inner nuclear layer; ONL, outer nuclear layer; PL, photoreceptor layer. * or ** indicates *p*<0.05 or *p*<0.01. Scale bar: 50 μm.

In order to further confirm the protective role of KITL in hereditary photoreceptor cell degeneration, we used an additional RP model, mice harboring a different mutant allele of *Pde6b*, *Pde6b*^*rd1*^ (hereafter called *rd1/rd1*) that was kept on a different background, FVB. The results were similar to those obtained with *rd10/rd10* mice (Supplemental Figure 8). Taken together, these results strongly indicate that overexpression of KITL prevents photoreceptor cell death and partially restores retinal function in both *rd10/rd10* and *rd1/rd1* mice.

## DISCUSSION

Here we present several lines of evidence that support the notion that activating KIT signaling can protect against both environmentally or genetically induced photoreceptor cell degeneration. First, KIT is expressed in mouse retina, including in its photoreceptor cells, and its activation leads to stimulation of MAPK and the PI3K/AKT pathways (Fig. 2H to J), confirming recent results by Too and collaborators (*55*). Second, photoreceptor cells respond to prolonged high-level light exposure by upregulating their endogenous KITL expression, which in turn activates KIT signaling (Fig. 1). Third, when KITL is experimentally overexpressed in *Kit+/+* eyes, it alleviates the effects of LD (Fig. 4). Fourth, when KIT signaling is genetically impaired, photoreceptors are more susceptible to LD, and this enhanced susceptibility cannot be overcome by additional experimental upregulation of KITL (Fig. 3 and Fig. 4H to J). Fifth, experimental overexpression of KITL partially protects against photoreceptor cell loss in *Kit+/+* mice harboring mutations in *Pde6b*, the mouse homologue of the gene mutated in human RP (Fig. 8 and Supplemental Figure 8).

The finding that in mice, KITL acts as a neuroprotective agent in retinal degenerative diseases may potentially lead to a therapeutic use of this factor in human retinal degenerative disorders for which therapeutic options are still limited. These options currently include gene and stem cell therapies, optogenetics, and retinal prostheses. Gene therapy has been successful for treating RPE65-associated Leber’s congential amaurosis, a retinal degenerative diseases caused by RPE65 mutations (*4, 8*). Nevertheless, gene therapy as well as the additional approaches mentioned above will always require major clinical and financial investments. In contrast, the application of an endogenous protective factor would seem to provide a more straightforward therapeutic avenue, with the added benefit that it may work regardless of the actual pathogenetic mechanism underlying photoreceptor degradation. In fact, AAV-mediated expression of neurotrophic factors or antioxidant genes such as RdCVF and NRF2 has proved beneficial in protecting photoreceptors (*16, 17, 56*).

Previous studies have indicated that Kit signaling can have both beneficial and deleterious effects (*30*). On the one hand, it is needed for cell development and survival, but on the other may also lead to formation of tumors, in particular gastrointestinal stromal tumors, adult acute myeloid leukemia, and melanoma (*30*). Hence, activation of Kit signaling should be used with caution in clinical applications. Nevertheless, local expression of KITL following intraocular injection of recombinant AAV might considerably reduce potential deleterious effects because gene expression is locally restricted and transfer of protein to other tissues is effectively blocked by the blood retinal barrier. In addition, the optimal choice of virus promoters and doses reduce the risk of virus-induced retinal damage (*57*). We observed that after 2 weeks of injection, high-level KITL expression was restricted to photoreceptor cells and the RPE (Fig. 4C and Supplemental Figure 4B), and that virus injection did not cause a significant change of retinal structure and cell death in adult mice over the duration of the experiment (Fig. 4D to G). Before initiating clinical studies, however, long term effects of virus injection should be evaluated and viral vectors might need optimization.

Although the underlying mechanisms of retinal degeneration are highly complex, oxidative stress may be common to many forms of retinal degeneration as well as to normal aging. Photoreceptor cells have a rich complement of mitochondria to accommodate the high metabolic rate and are more sensitive to oxidative stress than other cell types (*58*). The development of AMD, for instance, is tightly linked to oxidative damage (*1*), and antioxidant therapy can delay cell loss in retinal degenerative diseases and lead to reduced photoreceptor cell death in experimental animals (*59, 60*). Oxidative stress has also been observed in many other neurodegenerative conditions and antioxidant treatment is useful in their prevention (*25, 56, 60*). Not surprisingly, then, we found that LD enhanced expression levels of the endogenous antioxidative gene *Hmox1*, and this enhanced expression followed *Kitl* upregulation and depended on intact KIT signaling (Fig. 5). In addition, ectopic expression of HMOX1 in *Kit*^*Wps*^ mutant retinas (where *Hmox1* expression after LD is substantially reduced compared to *Kit+/+*) was able to protect these retinas at least partially against LD (Fig. 7). This function of Hmox1 was also observed in *Kit+/+* mice (*61*).

In line with previous observations (*62*), we found *Hmox1* expression to be regulated by nuclear accumulation of the transcription factor NRF2 (Fig. 6I and J), the master regulator of the antioxidant program. Interestingly, we found that under light stress, NRF2 accumulates in the nuclei particularly of photoreceptor cells (Fig. 6C and D) and is accompanied by upregulation of a series of antioxidant genes (Fig. 5 A). This process depends on Kit signaling as disruption of KIT signaling was able to prevent light-induced nuclear accumulation of NRF2 in photoreceptor cells (Fig. 6C and D, E-H). Moreover, it is known that the nuclear accumulation of NRF2 is regulated by AKT (*63*), a signaling molecule that lies downstream of KIT signaling, suggesting that KIT signaling regulates nuclear accumulation of NRF2 through AKT. Overexpression of Nrf2 effectively inhibits photoreceptor cell degeneration not only in light-induced retinal degeneration but also in mouse models of RP (*56, 64*). Collectively, these data support the notion that Kit signaling activates NRF2 and stimulates *Hmox1*expression to prevent photoreceptor cell degeneration.

In summary, our findings strongly provide evidence for KIT signaling in protecting photoreceptor cells against light-induced retinal damage as well as against inherited forms of retinal degeneration. We believe that these findings justify a detailed exploration of how this signaling pathway could be harnessed pharmacologically for prevention or delay of progressive photoreceptor cells and vision loss as seen in AMD, RP and other retinal degenerations. Hence, the findings have implications not only for our understanding of the relationship between genetic and environmental factors in retinal degeneration in general but also as a potential treatment for photoreceptor loss in humans.

## MATERIALS AND METHODS

### Mice and histology

All animal experiments were carried out in accordance with the approved guidelines of the Wenzhou Medical University Institutional Animal Care and Use Committee (Permit Number: WZMCOPT-090316). C57BL/6J mice were obtained from The Jackson Laboratory and *Kit*^*Wps*^ mice carrying a point mutation in the extracellular domain of KIT on a C57BL/6J background were as previously described (*46*). Because *Kit*^*Wps*^ homozygotes are infertile, crosses between *Kit*^*Wps*^ heterozygotes were set up to obtain *Kit*^*Wps/Wps*^ mice. To obtain *Kit*^*Wps/Wps*^ albino mice, we crossed *Kit*^*Wps*^ heterozygotes with BALB/c albino mice (*Tyr*^*c*^/*Tyr*^*c*^) to obtain *Kit*^*Wps/Wps*^;*Tyr*^*c*^/*Tyr*^*c*^ mice. For genotyping, the *Kit* gene was amplified with Kit-F and Kit-R, and the Rpe65 gene was amplified with Rpe65-F5 and Rpe65-R (Supplemental table 1) as previously described (*46, 48*). *rd1* (*Pde6b*^*rd1*^, FVB genetic background) and *rd10* (*Pde6b*^*rd10*^, C57BL/6J genetic background) mice were purchased from Vital River Laboratory (Beijing, China) and Jackson Laboratory. All mice were fed a standard laboratory chow and maintained at 21–23°C with a 12 h-light /12h-dark photoperiod.

Enucleated eyes from euthanized mice were fixed in 4% paraformaldehyde (PFA) overnight at RT and paraffin-embedded according to standard procedures. Paraffin sections (5 μm thickness) were prepared using a Leica microtome, deparaffinized by immersion in Xylene, and rehydrated. The sections were stained with hematoxylin for 5 minutes and eosin for 3 minutes, and mounted using resinene. The thickness of *Kit+/+* and *Kit*^*Wps/Wps*^ retinas was measured in the dorsal central region at a distance between 300 and 700μm from the optic nerve head.

### In vivo light injury and RNA-seq analysis

For light-induced retinal damage, 2-month-old albino mice were first dark-adapted for 12h and then exposed to LED light (15,000 lux) in non-reflective cages for the indicated number of days. For *Kit+/+* and *Kit*^*Wps/Wps*^ mice, 3-month-old mice were first dark-adapted for one day and then exposed to LED light (15,000 lux) in non-reflective cages for the indicated number of days. During light exposure, mice were allowed free access to food and water. For ERG measurements, mice were dark-adapted for one day after light exposure.

For RNA-seq analysis, total RNAs were extracted using TRIzol reagent (Invitrogen) after 1 day of light exposure following the time frame shown in Figure 1A. Purified RNA samples were used to generate RNA libraries for HiSeq 4000 (PE150 sequence).Experiments and data normalization were performed by Beijing Novogene (China).

### RT-PCR and quantitative RT-PCR

Total retinal RNAs were isolated from adult retinas using Trizol reagent (Invitrogen) according to the manufacturer’s protocol. Total RNAs for RT-PCR were converted to cDNA with a reverse transcriptase kit and random primers (Promega), according to the manufacturer’s manual (Promega). PCR products were size-fractionated by 2% agarose gel electrophoresis. For quantitative RT-PCR, the cDNA template was amplified (ABI PRISM 7500HT Sequence Detection System) using Power SYBR Green PCR Master Mix (4367659, ABI) under the following reaction conditions: 40 cycles of PCR (95°Cfor 15 s and 60°Cfor 1 min) after an initial denaturation (95°C for 2 min). The authenticity and size of the PCR products were confirmed using a melting curve analysis (using software provided by Applied Biosystems) and gel analysis. Sequences of primers used in this study are shown in Supplemental Tables 1 and 2.

### Electroretinography

Retinal function was evaluated by electroretinography (ERG) as previously described (*65*). Briefly, mice were dark-adapted overnight and anesthetized with a mixture of ketamine and xylazine. The pupils were dilated and the corneas were anesthetized with atropine sulfate and proparacaine hydrochloride. Then a silver loop electrode was placed over the cornea to record the ERGs, while needle reference and ground electrodes were inserted into the cheek and tail, respectively. Mice were stimulated by flash light varying in intensity from −5.0 to35 log scotopic candlepower-sec/m^2^ in a Ganzfeld dome (Roland Q400, Wiesbaden, Germany).For light-adapted ERG recordings, a background light of 30 cd/m^2^ was applied for 5-10 minutes to suppress rod responses. The stimulus light intensity was attenuated with neutral density filters (Kodak, Rochester, NY) and luminance was calibrated with an IL-1700 integrating radiometer/photometer (International Light, Newburyport, MA).

### Cell cultures, light damage, siRNA and transfection

The 661W photoreceptor cell line was generously provided by Dr. Muayyad Al-Ubaidi (Department of Cell Biology, University of Oklahoma Health Sciences Center, Oklahoma City, OK, USA). The cells were routinely cultured in Dulbecco’s modified Eagle’s medium (Gibco) supplemented with 10% FBS and 1% penicillin/streptomycin and kept at 37°C in a humidified atmosphere with 5% CO2.

For LD, 661W photoreceptor cells were kept at 37°C in a humidified atmosphere with 5% CO2 and exposed to high-intensity light (9000 lux) for 8 hours. Then the cells were subjected to cell count analysis and PI staining (1:200, Sigma).

For siRNA knockdown, specific siRNAs for mouse *Nrf2*, *Hmox* or *Kitl* and a negative control siRNA were designed and produced by Gene-Pharma (GenePharma, China). 20 nM si-*Nrf2*, si-*Hmox1* or si-*Kitl* was transfected into 661W cells at 40% confluency in culture dishes using LipoJet™ In Vitro Transfection Kit (SignaGen Laboratories) according to the manufacturer’s protocol. An equivalent amount of control siRNA (si–C) was used as a negative control. The sequences for all siRNAs are shown in Supplementary Table 3.

For DNA transfections, plasmids of pCDNA3.1-encoding either KIT^WT^cDNA or KIT^Wps^ cDNA were from Dr. Xiang Gao (Nanjing University). 661W cells were seeded in 6-well plates at 7.5 × 10^4^ cells/well before transfection. When the cell density reached 30% confluence, cells were transfected with appropriate dilutions of 2 μg plasmid DNA using PolyJet™ reagent (SignaGen) at the indicated concentrations following the company’s instructions.

### Immunohistochemistry and TUNEL assay

Information of antibodies used in immunofluorescence (IF) is shown in Supplemental Table 4. For immunoassays of KITL, KIT and HMOX1, eyes were freshly embedded in OCT compound and snap frozen. Cryosections (14 μm) were post-fixed in ice-cold acetone for 15 minutes and permeabilized with 0.1% Triton-X-100 for 10 minutes. For blocking samples, 3% BSA was used with incubation at room temperature (RT) for 30 minutes. Rat anti-KIT (ACK45), goat anti-KITL and rabbit anti-HMOX1 antibodies were used at 4°C overnight and Alexa 488-conjugated donkey anti-rat, anti-goat or anti-rabbit antibodies were used at RT for 1 hour. The sections were examined and photographed with a Zeiss fluorescence microscope.

For other immunoassays, the corneas were penetrated with a needle in 1×PBS and the eyes fixed in 4% PFA for 1 hour at RT. The sections (5μm) were heated in boiled citrate buffer for 2 min for epitope retrieval. For blocking samples, 5% BSA was used at RT for 1 hour. Primary antibodies were: mouse anti-rhodopsin, rabbit anti-opsin, mouse anti-PKCα, rabbit anti-OTX2, goat anti-KITL, rabbit anti-KIT (AB5506), rabbit anti NRF2 and rabbit anti-GFAP. The antibodies were all used at RT for 2 hours. The primary antibodies were revealed with Alexa 594-conjugated donkey anti-mouse or anti-rabbit and Alexa 488-conjugated donkey anti-goat or mouse antibodies at RT for 1 hour.

Cultured cells were fixed in 4% PFA for 25 min at RT and permeabilized with 0.1% Triton X-100. 5% BSA was used to block the samples at RT for 30 min. Rat anti-KIT (ACK2) antibody was used at RT for 2 hours.

For TUNEL assays, in situ cell death detection kit (Roche, Mannheim, Germany) was used according to the manufacturer’s protocol.

### Western blots and assays of protein phosphorylation

For analysis of phosphorylation of KIT, mice were anesthetized with a mixture of ketamine and xylazine, and 1μl of KITL (5.6 nM) was intravitreally injected into the eye. One hour thereafter, retinas were isolated and subjected to western blot analysis. For protein extraction, retinas were isolated in DMEM and transferred to 1.5 ml centrifuge tubes containing a mixture (100μl) of membrane protein lysis solution (Byotime, China) and protease inhibitors (Cocktail Set I, Calbiochem). Retinas were lysed on ice for 10 minutes with a Micro Tissue Grinder. Samples were separated by 8% SDS-PAGE and transferred onto PVDF membranes (Millipore, Molsheim, France). After blocking the membranes with 5% non-fat milk at RT for 2 hours, rat anti-KIT and rabbit anti-p-KIT antibodies (Tyr 567/569) were used and incubated at 4°C overnight. Goat anti-rabbit IgG (H+L) IRDye® 800CW and goat anti-rat IgG (H+L) IRDye® 800CW antibodies were used and incubated at RT for 1 hour in the dark. The protein bands were scanned using a LI-COR machine. Mouse anti-α-tubulin or β-actin antibodies were used as internal controls to normalize for the amount of total proteins. Primary antibodies were: Rabbit anti-MDA, goat anti-KITL, rabbit anti-HMOX1, rabbit anti-ERK, rabbit anti-p-ERK, rabbit anti-AKT and rabbit anti-p-AKT. Information of antibodies used in western blot is shown in Supplemental Table 3. These antibodies were used at 4°C overnight. The quantification of the protein bands was done using the software of Bandscan.

### AAV8 vector construction and virus injection

The vector of pcDNA3.0-*Kitl* was a gift from Dr. Bernhard Wehrle-Haller (University of Geneva, Switzerland). The coding sequence of the mouse *Kitl* was cloned from the vector and inserted into the AAV-MCS8 empty vector (Genechem, China). The coding sequence of the mouse *Hmox1* gene was amplified by reverse transcription PCR using the primers for Hmox1-F and Hmox1-R as shown in Supplemental Table S1. The full-length cDNA of *Hmox1* was cloned from *Kit+/+* retinal total cDNA and inserted into the AAV-MCS8 empty vector (Genechem, China) via *Age*I restriction sites introduced by the PCR reaction. The AAV8-GFP vector was purchased from Genechem Company (Shanghai, China).

Virus was injected into the subretinal space as previously described (*56*). Approximately 0.5 μl of AAV8 supernatant (2.2 × 10^12^ genome copies/ml) was introduced into the subretinal space (of 2-3 month old mice) using a pulled angled glass pipette under direct observation aided by a dissecting microscope under dim light. Following injections, 1% atropine eye drops, tetracycline, and cortisone acetate eye ointments were applied. For subretinal injection, neonatal *rd1* and *rd10* mice were kept on ice for 2 minutes and then subretinally injected with approximately 0.3 μl of AAV8 using micro-injection glass pipettes.

### Statistical analysis

Data are from at least three replicates for each experimental condition and are represented as mean±standard error of the mean (SEM). Two-way ANOVA was used to determine the significance of differences between population means. *P*<0.05 was considered to be statistically significant. Significant differences between groups are noted by *or **.

## Acknowledgments

We thank Drs. Xiang Gao and Wei Li for providing the *Kit*^*Wps*^ mice and reagents, Dr. Bernie Wehrle-Haller for providing reagents, Drs. Muayyad R. Al-Ubaidi and Feng Gu for 661W cell line, and Drs. Heinz Arnheiter, Wenjun Xiong, and Wei Li (NEI), and for thoughtful comments and editing of the manuscript.

## Funding

We gratefully acknowledge the National Natural Science Foundation of China (81800838, 81770946, 81600748, 81870664), the Zhejiang Provincial Natural Science Foundation (LY18H120007, LY13C090004, LZ12C12001), and the Research Development Grant of Wenzhou Medical University for generous funding. This study was also supported by the Project of State Key Laboratory of Ophthalmology, Optomertry and Visual Science, Wenzhou Medical University (437201804G).

## Author Contributions

Conceived and designed the experiments: H Li, L Hou. Performed the experiments: H Li, L Lian, B Liu, Y Chen, J Yang, S Jian, J Zhou. Analyzed the data: H Li, L Hou. Contributed reagents/materials/analysis tools: H Li, Y Xu, X Ma, J Qu, L Hou. Wrote the manuscript: H Li, L Hou.

## Competing interests

The authors declare no competing financial interests.

