## Supplementary material for "KIT ligand protects against both light-induced and genetic photoreceptor degeneration": Suppl Materials

**Supplemental Materials**

**Supplemental Table 1. Sequences of primers used for RT-PCR**

| Primer names | Sequences |
|---|---|
| Kit-g-F | TTGACCCAGAGAAAGCTGTG |
| Kit-g-R | GAGAACAGGGAAGCAACTACC |
| Hmox1-c-F | GGCATGCCTTCCGCATACAACCAGTG |
| Hmox1-c-R | AGGCGCGCCTTCAGGTATCTCCCTCCATT |
| Rpe65-g-F | CTGACAAGCTCTGTAAG |
| Rpe65-g-R | CATTACCATCATCTTCTTCCA |

**Supplemental Table 2. Sequences of primers used for gene expression**

| Gene name | NCBI Gene ID | Forward primer | Reverse primer |
|---|---|---|---|
| Gapdh | 14433 | ACCACAGTCCATGCCATCAC | TCCACCACCCTGTTGCTGTA |
| Lif | 16878 | AACGATGGTGTCACCCTGAAG | TATATACTGGAGCCGTGGTCT |
| Relb | 19698 | TAGCCTTGCACCGCTTGC | CAATTCATCTGTGGTCCTGGAG AC |
| Epha2 | 13836 | GGTTCTCACCCAACTTCCAT | CGAAGTCCAACAAAACAACTT CCT |
| c-Fos | 14281 | CAGAGCGGGAATGGTGAAGA | CTGTCTCCGCTTGGAGTGTA |
| Nfkb1 | 18033 | GGTCACCCATGGCACCATAA | AGCTGCAGAGCCTTCTCAAG |
| Ngf | 18049 | GGAGCGCATCGAGTGACTT | CCTCACTGCGGCCAGTATAG |
| Myd88 | 17874 | TGTTCTTGAACCCTCGGACG | TTCTGGCAGTCCTCCTCGAT |
| Hmox1 | 15368 | CAGAAGAGGCTAAGACCGCC | GCAGTATCTTGCACCAGGCTA |
| Nrf2 | 18024 | CATGAGACATCTGCTCGCCT | ACTGCCCAAGCAGAGTTAGG |
| Kit | 16590 | AAGCGTCTCCACCATTCA | GGAGCGGTCAACAAGGAA |
| Kitl-V1 | 17311 | TTATGTTACCCCCTGTTGCAG | CTGCCCTTGTAAGACTTGACTG |
| Kitl-V2 | 17311 | TCCCGAGAAAGGGAAAGC | CTGCCCTTGTAAGACTTGACTG |

**Supplemental Table 3. Sequences of siRNAs used in the study.**

| siRNA names | siRNA Sequences |
|---|---|
| si-Nrf2-1 | GCAGGACAUGGAUUUGAUUTT / AAUCAAAUCCAUGUCCUGCTT |
| si-Nrf-2 | CCGAAUUACAGUGUCUUAATT / UUAAGACACUGUAAUUCGGTT |
| si-Kitl-1 | GCUCCUAUUUAAUCCUCUUTT / AAGAGGAUUAAAUAGGAGCTT |
| si-Kitl-2 | GCUACGAGAUAUGGUAAUATT / UAUUACCAUAUCUCGUAGCTT |
| si-Hmox1-1 | CCACCAAGGAGGUACACAUT / AUGUGUACCUCCUUGGUGGTT |
| si-Hmox1-2 | GCUGACAGAGGAACACAAATT / UUUGUGUUCCUCUGUCAGCTT |
| si-NC | UUCUCCGAACGUGUCACGUTT / ACGUGACACGUUCGGAGAATT |

**Supplemental Table 4. Antibodies used in immunofluoresence (IF) and western blot (WB).**

| Antibodies | Source | Cat# | Dilution |
|---|---|---|---|
| Rabbit polyclonal to GFAP | Abcam | ab7260 | 1:500 (IF) |
| Rabbit polyclonal to KIT | Abcam | ab5506 | 1:100 (IF)<br>1:800(WB) |
| Rabbit polyclonal to NRF2 | Abcam | AB137550 | 1:200 (IF)<br>1:1000(WB) |
| Mouse monoclonal to Rhodopsin | Millipore | MABN15 | 1:200 (IF) |
| Rabbit polyclonal to Opsin | Millipore | AB5745 | 1:200 (IF) |
| Rabbit polyclonal to OTX2 | Millipore | AB9566 | 1:200 (IF) |
| Mouse monoclonal to PKCα | Santa Cruz Biotechnology | sc-8393 | 1:50 (IF) |
| Rabbit polyclonal to HMOX1 | Santa Cruz Biotechnology | sc-10789 | 1:50 (IF)<br>1:200(WB) |
| Rabbit polyclonal to p-KIT (Tyr568/570) | Santa Cruz Biotechnology | sc-18076-R | 1:1000(WB) |
| Mouse monoclonal to α-Tubulin | Santa Cruz Biotechnology | sc-53646 | 1:1000(WB) |
| Mouse monoclonal to β-Actin | Santa Cruz Biotechnology | sc-130300 | 1:1000(WB) |
| IRDye® 800CW Donkey anti-mouse lgG(H+L) | LI-COR | C70502-02 | 1:5000(WB) |
| IRDye® 680RD Donkey anti-rabbit lgG(H+L) | LI-COR | C30420-01 | 1:5000(WB) |
| IRDye® 800CW Donkey anti-goat lgG(H+L) | LI-COR | C11117-05 | 1:5000(WB) |
| IRDye® 800CW Goat anti-rat lgG(H+L) | LI-COR | C11024-01 | 1:5000(WB) |
| IRDye® 680RD Goat anti-mouse lgG(H+L) | LI-COR | C30502-01 | 1:5000(WB) |
| Rat monoclonal to KIT (ACK45) | BD Pharmingen | 553868 | 1:50 (IF)<br>1:200(WB) |
| Rat monoclonal to KIT (ACK2 / CD117) | Invitrogen | 14-1172-82 | 1:50(IF) |
| Alexa 594-conjugated donkey anti-rabbit lgG(H+L) | Invitrogen | 1827674 | 1:500(IF) |
| Alexa 488-conjugated donkey anti-mouse lgG(H+L) | Invitrogen | 1796361 | 1:200(IF) |
| Alexa 594-conjugated donkey anti-mouse lgG(H+L) | Invitrogen | 1820027 | 1:500(IF) |
| Alexa 488-conjugated donkey anti-rabbit lgG(H+L) | Invitrogen | 1834802 | 1:200(IF) |

| | | | |
|---|---|---|---|
| Alexa 488-conjugated donkey anti-goat lgG(H+L) | Invitrogen | 1605893 | 1:200(IF) |
| Goat polyclonal to KITL/SCF | R&D systems | 455-MC | 1:50(IF)<br>1:800(WB) |
| Rabbit polyclonal to p44/42 MAPK (Erk1/2) | Cell signaling technology | 4695S | 1:2000(WB) |
| Rabbit polyclonal to p-p44/42 MAPK (Erk1/2) (Thr202/Tyr204) | Cell signaling technology | 4370S | 1:2000(WB) |
| Rabbit polyclonal to AKT | Cell signaling technology | 9272S | 1:2000(WB) |
| Rabbit polyclonal to p-AKT (Ser473) | Cell signaling technology | 4060S | 1:2000(WB) |

### Supplement figures

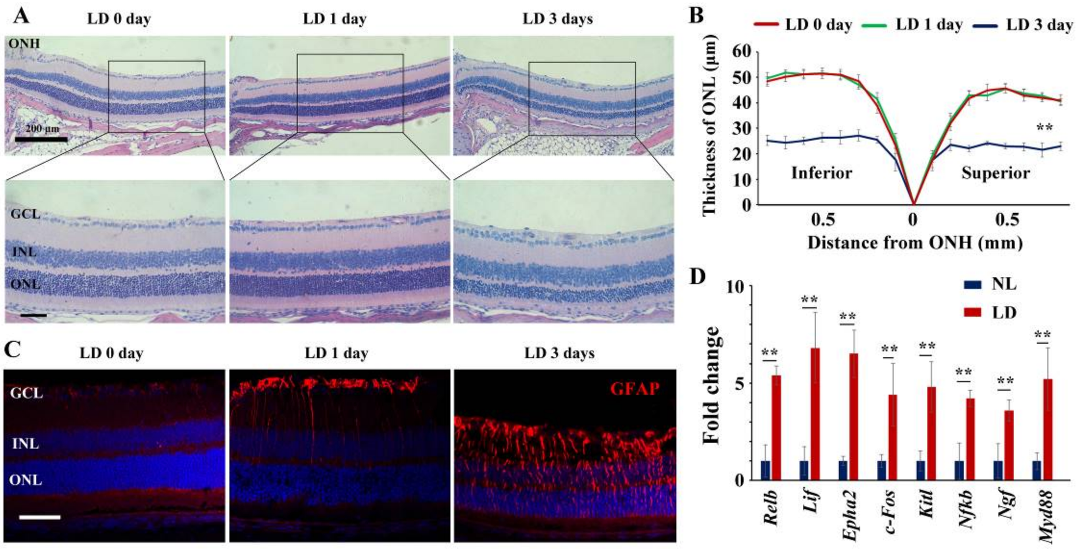


**Supplemental Figure 1. High intensity light induces retinal damage in albino mice.** (**A**, **B**) Histological images of HE staining from the retinas (**A**) and the curve diagram shows the thickness of the outer nuclear layer (ONL) based on the results of hematoxylin-eosin staining (**B**). Note that the structure was significantly changed in retinas treated with high intensity LD for 3 days instead of retinas treated with LD for 1 day. (**C**) Immunofluorescence images of anti-GFAP antibody in retinas from 2-month-old albino mice kept under the indicated condition. Note that LD induces increased expression of GFAP in the retinas treated for 1 day or 3 days. (**D**) RT-PCR data show changes in transcript levels of MAPK pathway genes in albino retinas in response to LD. GCL, ganglion cell layer; INL, inner nuclear layer; IS, inner segments; ONL, outer nuclear layer; OS, outer segments；PL, photoreceptor cell layer;. ** indicates $p$<0.01. Bar: 50 µm.

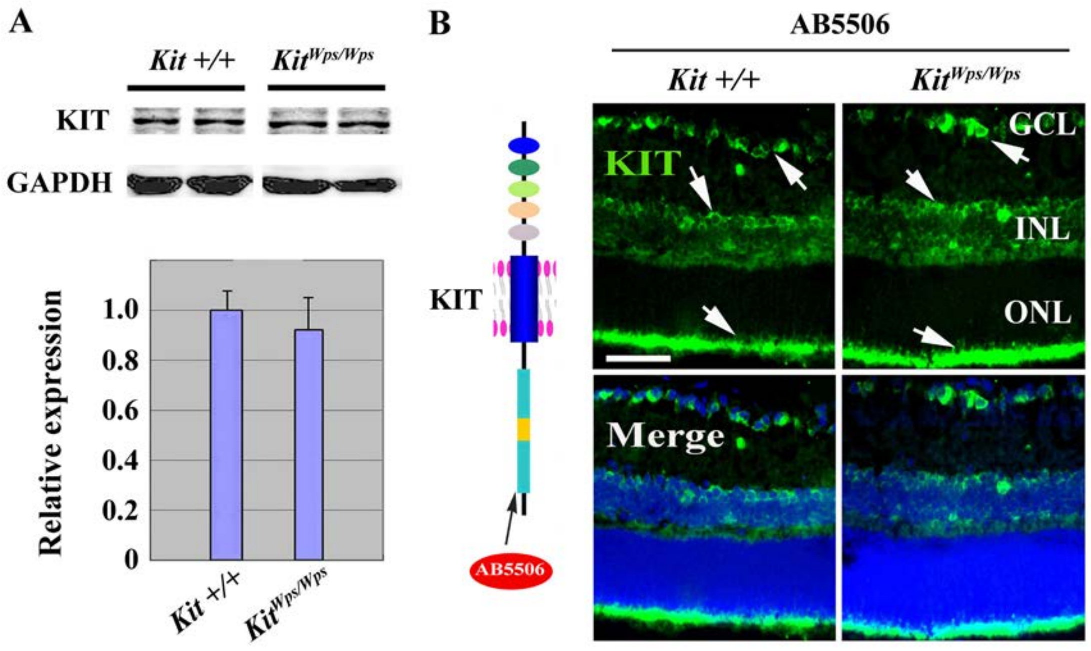


**Supplemental Figure 2. The *Kit*$^{Wps}$ mutation does not affect KIT expression.** (**A**) Western blots show expression of KIT in *Kit*+/+ and *Kit*$^{Wps/Wps}$ retinas (upper panels) using antibodies reacting with the KIT C terminus (AB5506) and bar graph shows no apparent difference of KIT expression between wildtype and mutant retinas (lower panels). (**B**) Schematic representation of KIT protein structure and the location recognized by the AB5506 antibody (left panels) and immunostaining images of AB5506 antibody in 3-month-old *Kit*+/+ and *Kit*$^{Wps/Wps}$ retinas (right panels). GCL, ganglion cell layer; INL, inner nuclear layer; ONL, outer nuclear layer. Bar: 50 µm.

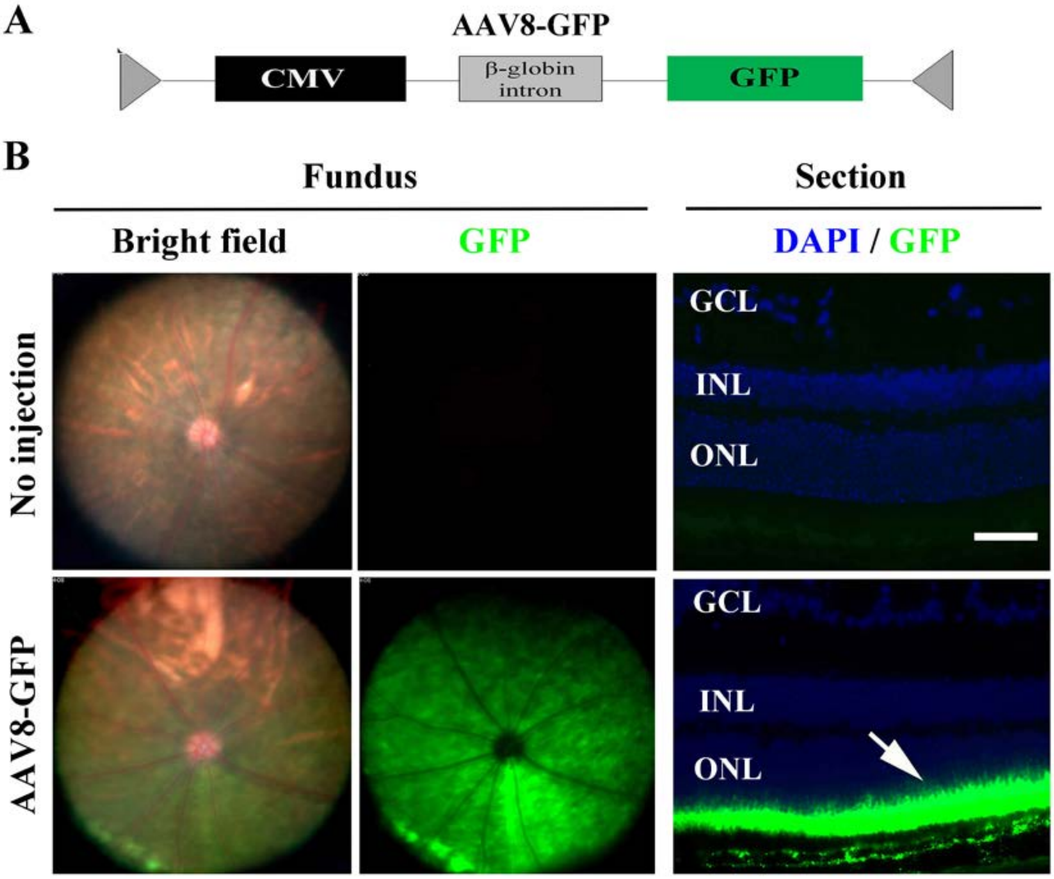


**Supplemental Figure 3. AAV8-GFP virus specifically infects photoreceptor cells throughout the *Kit*$^{Wps}$ mutant retina.** (**A**) Schematics of the AAV8-*GFP* construct. (**B**) Bright field and GFP fluorescence fundus photographs after AAV8-GFP infection (left panels) and a representative cross-section image of an AAV-GFP–infected retina (right panels). The ONL shows a high degree of infection (green). ONL, outer nuclear layer. Bar: 50 µm.

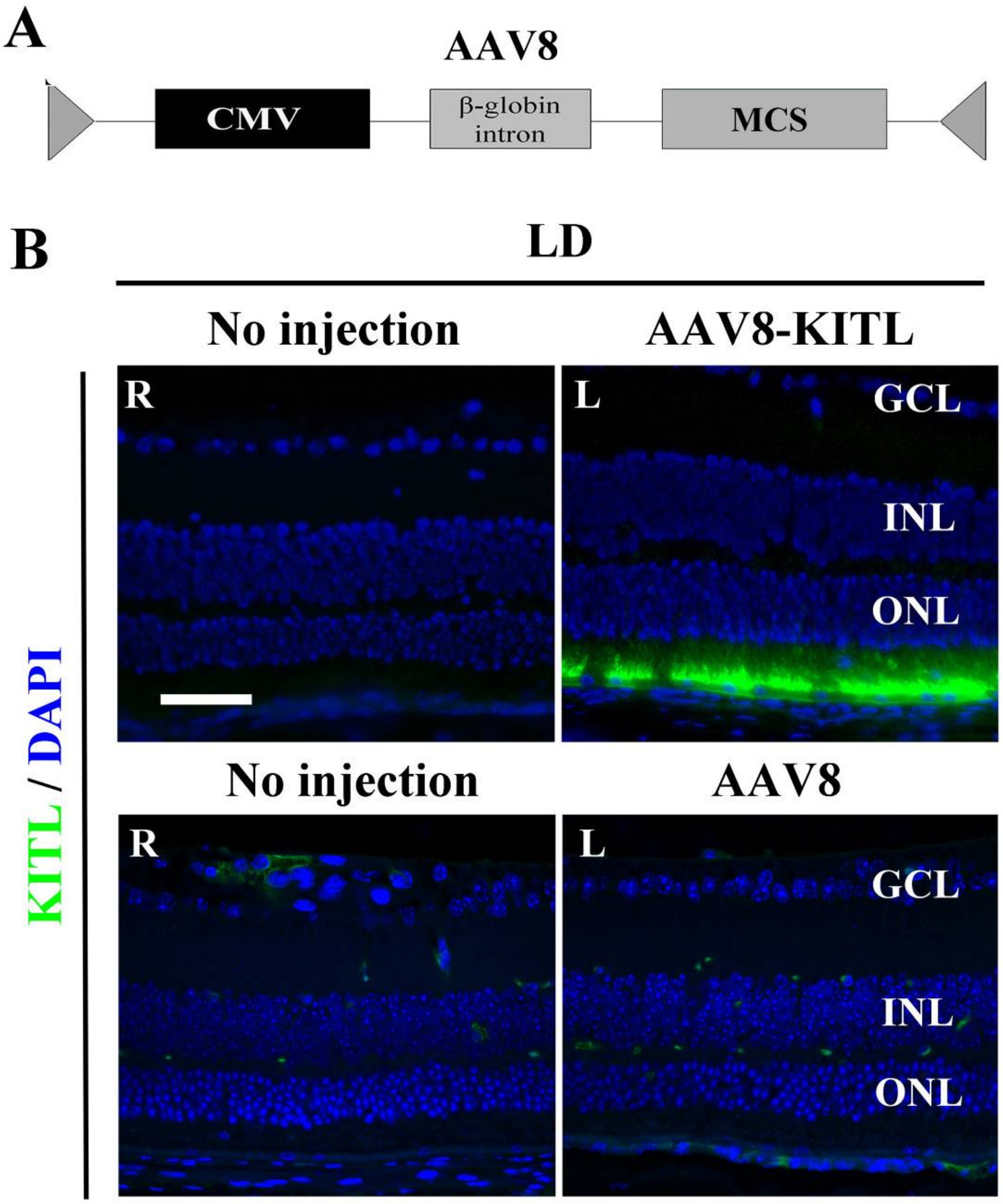


**Supplemental Figure 4. High expression of KITL in the retina infected by AAV8-KITL virus after LD treatment. (A**) Schematic depiction of the AAV8 construct. (**B**) Immunofluorescence images of anti-KITL in the retinas under the indicated conditions. Right (R) eye and left (L) eye are from the same mouse, whereby (R) has not been injected (both upper and lower panel) while (L) in the upper panel has been injected with AAV8-KITL and (L) in the lower panel has been injected with control AAV8. GCL, ganglion cell layer; INL, inner nuclear layer; ONL, outer nuclear layer. Bar: 50 μm.

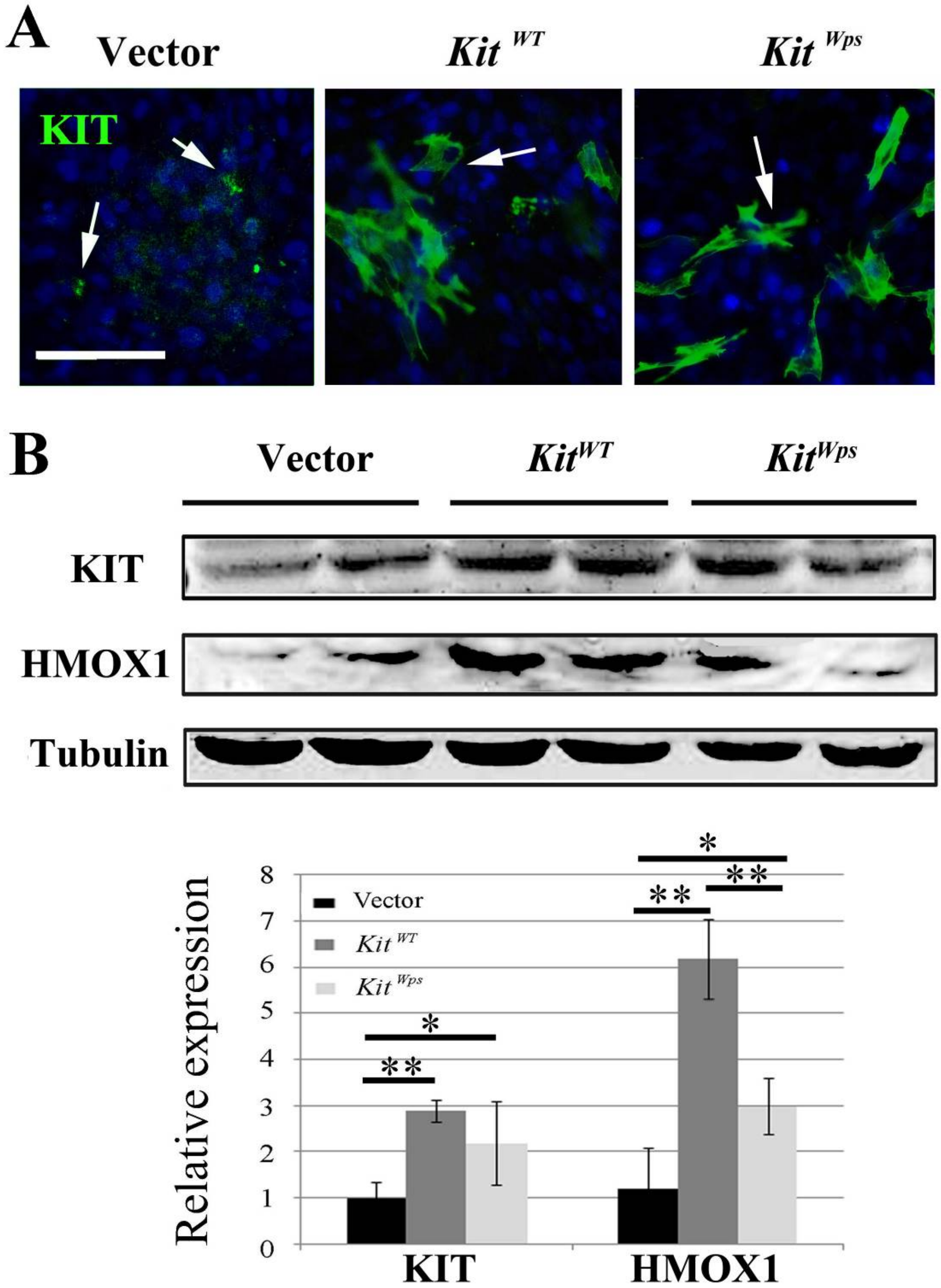


**Supplemental Figure 5. Overexpression of KIT upregulates *Hmox1* expression in 661W photoreceptor cells.** (**A**) Immunostaining using a KIT antibody (ACK2 / CD117) after transfection of $Kit^{WT}$ (wildtype Kit cDNA plasmid) or $Kit^{Wps}$ (Kit-Wps mutant cDNA plasmid) in 661W cells with 2.8 nM of KITL. (**B**) Immunoblotting of KIT and HMOX1 expression in 661W cells transfected with $Kit^{WT}$ or $Kit^{Wps}$ plasmid (upper panels), and quantification of the immunoblotting results (lower panel). Note that overexpression of wildtype Kit but not $Kit^{Wps}$ led to upregulation of *Hmox1* expression. * or ** indicates *p*<0.05 or *p*<0.01. Bar: 50 μm.

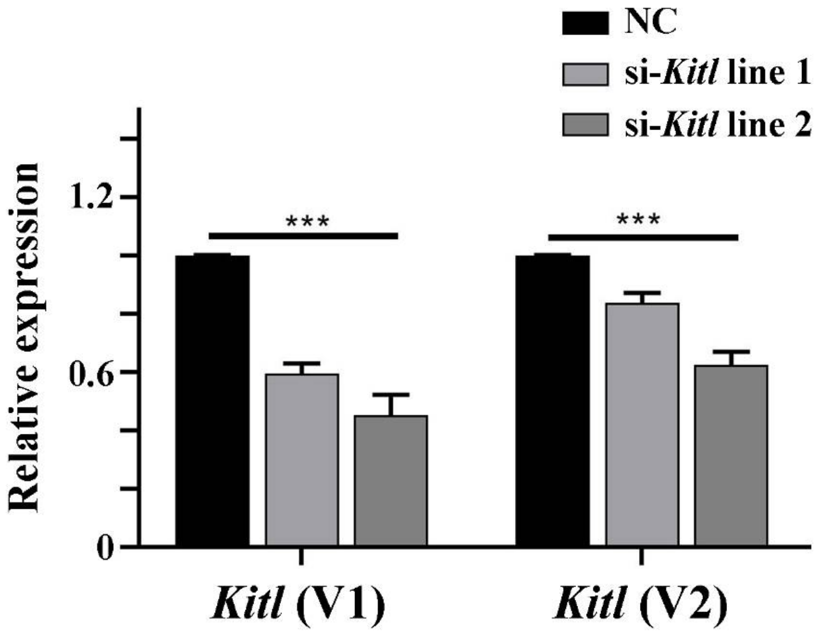


**Supplemental Figure 6. Messenger RNA expression levels of *Kitl* are reduced after transfection with Kitl-si-RNAs in 661W cells.** The transcript variant 1 (V1) contains the full length Kitl mRNA sequence while the transcript variant 2 (V2) corresponds to a Kitl mRNA lacking exon 6.

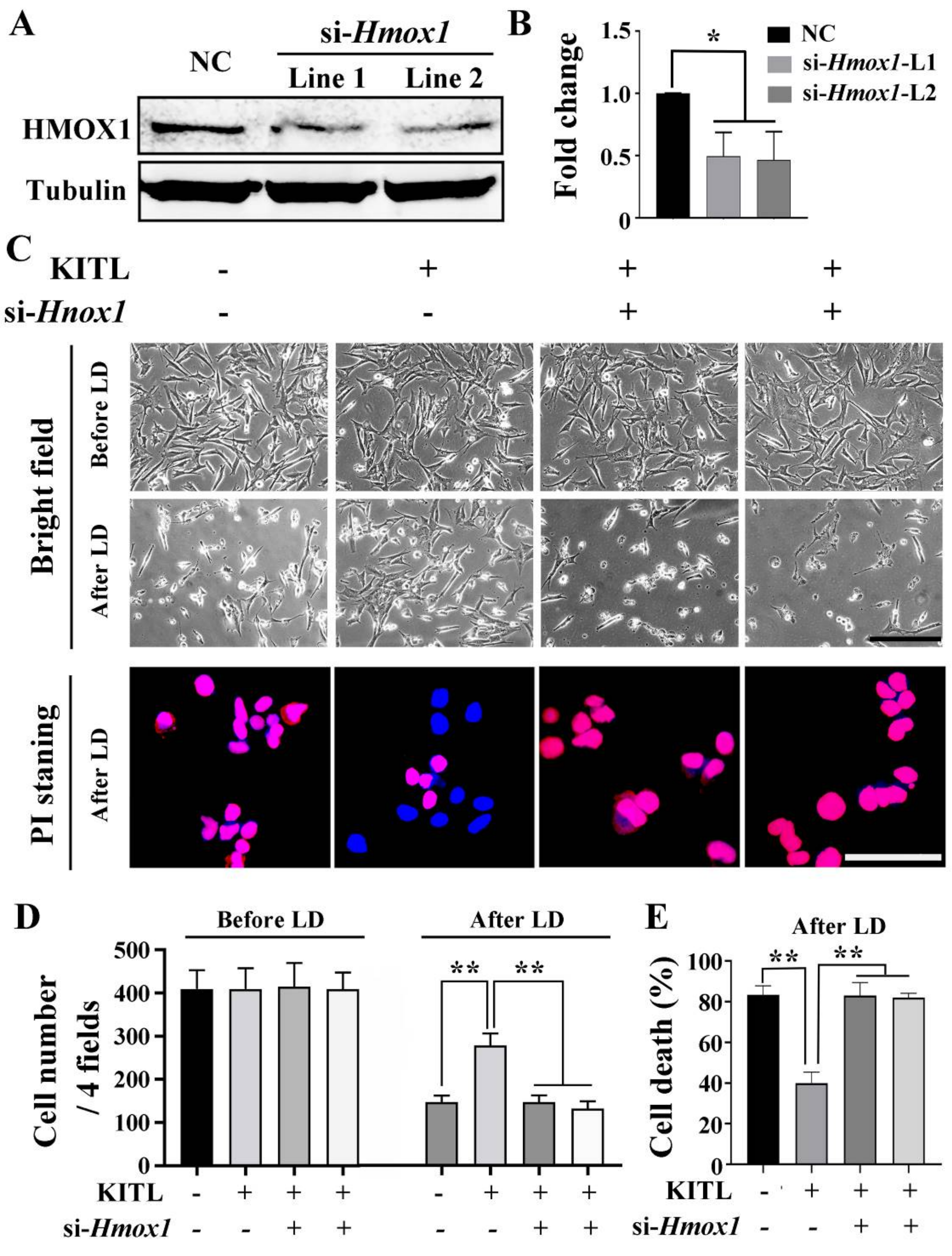


**Supplemental Figure 7. Protective role of KITL depends on HMOX1 in** light-induced photoreceptor cell death. **(A)** Western blot analysis of HMOX1 in 661W photoreceptor cells after si-RNA treatment. (**B**) Quantification of western blot bands shows expression levels of HMOX1. Note that both si-*Hmox1* lines show reduced expression of HMOX1. (**C**) Bright field images of photoreceptor cells under the indicated conditions (upper panels) and Propidium Iodide (PI) staining of photoreceptor cells after LD (down panels). (**D, E**) Bar graphs show the cell numbers before and after high-intensity light treatment (**D**) and the numbers of PI positive cells after LD treatment (**E**). ** indicates *p*<0.01. Scale bar, 20 μm.

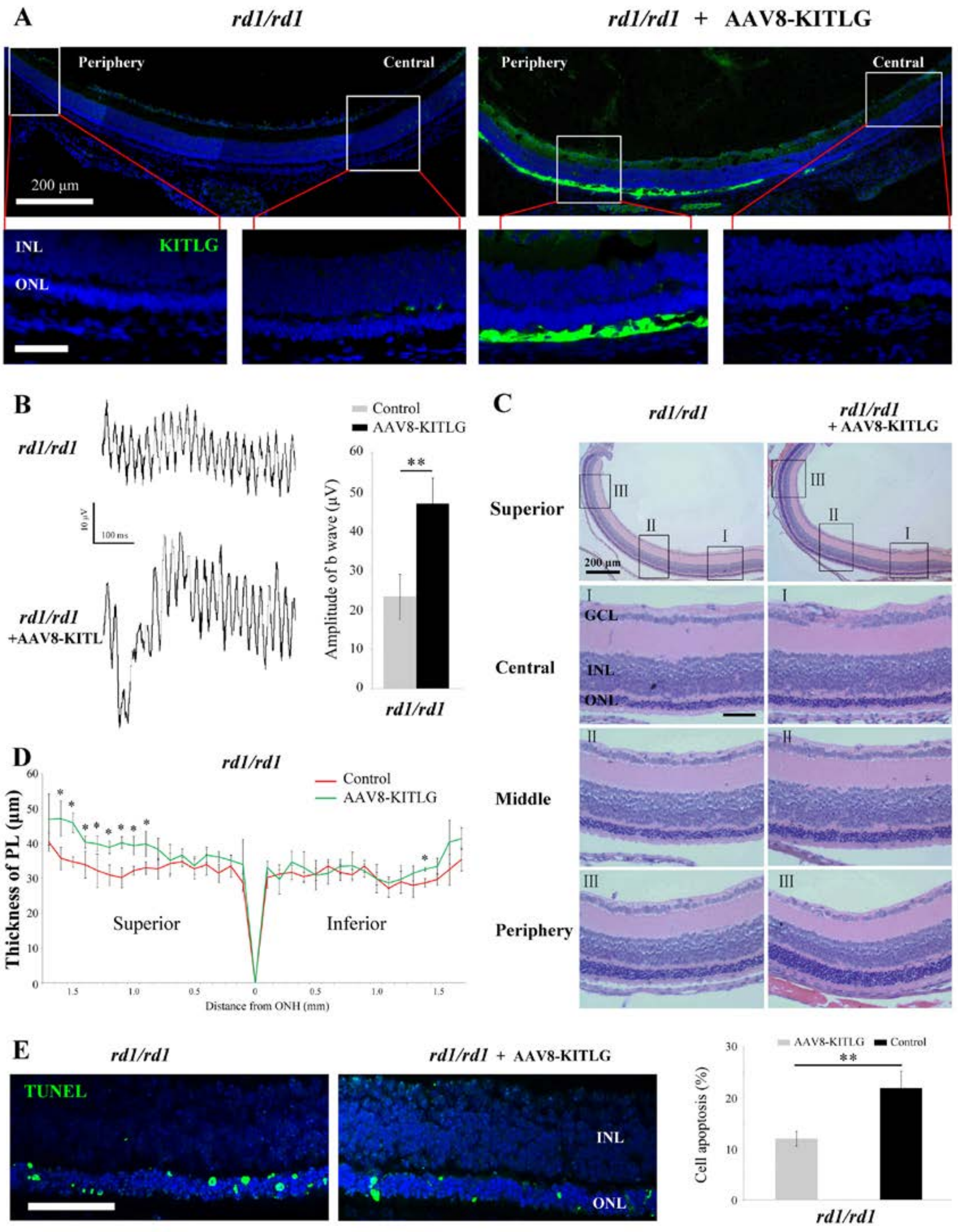


**Supplemental Figure 8. Ectopic expression of KITL prevents retinal degeneration of *rd1* homozygous mice.** (**A**) Expression of KITL in *rd1/rd1* retinas

2 weeks after infection with AAV8-KITL virus, or AAV8 (control) infection, was visualized by immunofluorescence. Note that the subretinal injection was performed in *rd1/rd1* mice at postnatal day 1. (**B**) ERG scotopic traces obtained from *rd1/rd1* mice with or without infection of AAV8-KITL virus (n=5) and kept under normal environmental condition for 20 days (right panels). Bar graph shows the quantification of the amplitude of the b-wave from standard response based on the results of the ERG scotopic traces (left panels). (**C** and **D**) Histological analysis of *rd1/rd1* retinas infected with AAV8-KITL virus (n=5) or control (n=5) (**C**) and the curve diagram showing the thickness of total ONL from *rd1/rd1* retina under the indicated conditions (**D**). (**E**) Apoptosis of photoreceptor cells from *rd1/rd1* mice with AAV8-KITL infection or control was analyzed by TUNEL assay and presented as fluorescent images (left panels) and as bar graphs after quantitation (right panel). GCL, ganglion cell layer; INL, inner nuclear layer; ONL, outer nuclear layer; PL, photoreceptor cell layer. * or ** indicates $p<0.05$ or $p<0.01$. Bar: 50 μm.
